# Positive feedback induces switch between distributive and processive phosphorylation of Hog1

**DOI:** 10.1101/2020.05.05.078352

**Authors:** Maximilian Mosbacher, Sung Sik Lee, Matthias Peter, Manfred Claassen

## Abstract

Cellular decision making often builds on ultrasensitive MAPK pathways. The phosphorylation mechanism of MAP kinase has so far been described as either distributive or processive, with distributive mechanisms generating ultrasensitivity in theoretical analyses. However, the *in vivo* mechanism of MAP kinase phosphorylation and its regulation by feedback loops remain unclear. We thus characterized the regulation of the MAP kinase Hog1 in *Saccharomyces cerevisiae*, which is transiently activated in response to hyperosmolarity. Specifically, we combined Hog1 activation data from different modalities and multiple conditions. We constructed ODE models with different pathway topologies, which were then assessed *via* parameter estimation and model selection. Interestingly, our best fitting model switches between distributive and processive phosphorylation behavior via a positive feedback loop targeting the MAP kinase-kinase Pbs2. Simulations further suggest that this mixed mechanism is required not only for full sensitivity to stimuli, but also to ensure robustness to different perturbations.

## Introduction

Pathways integrating extracellular inputs often display an ultrasensitive response, in which beyond an input threshold, small changes in the input lead to large changes in the output. This behavior results in an essentially binary response, acting as a switch in the overall signaling cascade (Figure 1A) (Altszyler et al., 2017). Ultrasensitivity has been experimentally observed in various signaling systems and plays an important role in cellular decision making (Ferrell and Ha, 2014). Theoretical studies suggest that multi-tiered multisite phosphorylation cascades are inherently able to create ultrasensitivity (Huang and Ferrell, 1996), with even single multisite phosphorylation resulting in ultrasensitivity and bistability (Markevich et al., 2004). In particular, the specific type of phosphorylation mechanism can alter signal response dynamics (Salazar and Hofer, 2009). For dual phosphorylation, two distinct mechanisms are recognized (Figure 1B). A distributive mechanism involves two consecutive reaction events, with kinase and substrate dissociating after each phosphorylation step, while for a processive mechanism, two phosphorylation reactions are induced in a single concerted reaction event (Salazar and Hofer, 2009). Dual phosphorylation is a particularly widespread mechanism involved in activating mitogen-activated protein (MAP) kinases, which regulate the cellular responses to many intra- and extracellular signals. However, investigating the impact of different phosphorylation mechanisms is challenging and normally relies on mathematical models integrating typically difficult-to-measure temporal dynamics of specific protein species. The majority of theoretical studies have reduced assumptions to *“all-or-none”* conditions – either distributive or processive – with distributive mechanisms correlating with ultrasensitivity while processive mechanisms associate with a graded, non-ultrasensitive response (Patwardhan and Miller, 2007). Thus to date, theoretical studies generally suggest that ultrasensitive kinase phosphorylation events *in vivo* should be governed by a distributive phosphorylation mechanism (Huang and Ferrell, 1996; O’Shaughnessy et al., 2011), and some experimental data from mammalian cells support this hypothesis (Burack and Sturgill, 1997; Ferrell and Bhatt, 1997). However, in more complex cases of multisite phosphorylation with more than two phospho-sites, behaviors have been observed that are not well explained by either a processive or distributive mechanism (Jeffery et al., 2001). Some of these multisite phosphorylation events involve different kinases with distinct kinetic properties, which may complicate the analysis (Koivomagi et al., 2011). Altogether, understanding kinase phosphorylation mechanisms governing ultrasensitive responses has proven particularly challenging in cases of multisite phosphorylation and other feedback regulation, and requires additional studies to explore the theoretical potential and experimental validation of mixed phosphorylation mechanism.

**Figure 1:**
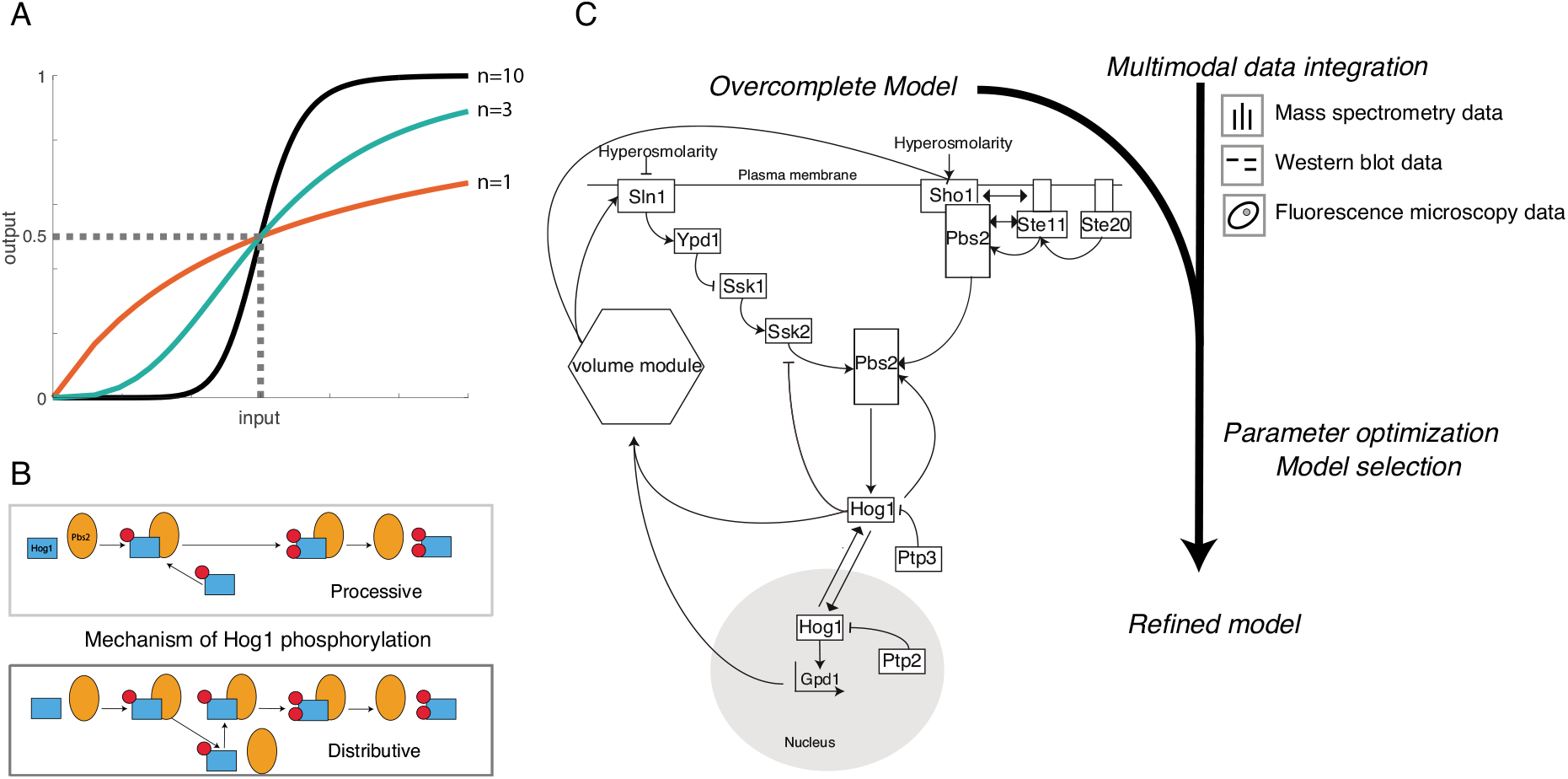
Generation of an overcomplete model of the Hog1 pathway and parameter optimization. (A) A Hill function quantifies highly ultrasensitive (black line, Hill coefficient = 10), mildly ultrasensitive (green line, Hill coefficient = 3), or strictly Michaelian (red line, Hill coefficient = 1) input-output behavior. EC50 value (dashed line) indicates the input strength at which 50% of the maximal output is reached. (B) The choice of Hog1 phosphorylation mechanism has a significant impact on the resulting input-output behavior and can be mainly processive or distributive. For the processive mechanism, the second phosphorylation step immediately follows the first without Hog1 disassociating from Pbs2. For the distributive mechanism Hog1 disassociates from Pbs2 after every phosphorylation step and needs to re-bind for further phosphorylation to take place. (C) Simplified scheme of the workflow to distinguish between different Hog1 phosphorylation mechanisms. The overcomplete ODE model allows enumeration of varying model topologies that differ in negative and positive feedback mechanisms, and Hog1 phosphorylation mechanism. Multimodal data was collected from literature or was newly self-generated, including fluorescence microscopy, mass spectrometry and western blot experiments. The data reflects various experimental conditions such as different salt concentrations and pulse frequencies, deletion mutants and kinase inhibition. Data sets were used to optimize the parameter values of the different submodels of the overcomplete model thus giving rise to refined models from which the best fitting was selected

To address these questions, we choose to investigate the mechanisms conferring ultrasensitivity in the <u>H</u>igh <u>O</u>smolarity <u>G</u>lycerol (HOG) pathway in *Saccharomyces cerevisiae,* a well-studied MAP-kinase pathway which requires dual phosphorylation of the MAP kinase Hog1. Hog1 activation is needed to re-establish the balance between internal and external pressures upon osmotic shock. Upon exposure of cells to high osmolarity conditions, the two membrane-localized osmo-sensors Sho1 and Sln1 activate either the MAPKKKs Ste11 or Ssk2,22, which converge on the MAPKK Pbs2 (Figure 1C). Activated Sho1 recruits Pbs2, which acts both as a MAPKK to phosphorylate the MAP kinase Hog1 and as a scaffold recruiting other upstream kinases including its own activator Ste11. Additionally, Ste50 and the three transmembrane proteins Msb2, Hrk1 and Opy2 are needed for full Ste11 activation by recruiting various co-stimulators to the cell membrane (Ekiel et al., 2009; Tanaka et al., 2014). The partially redundant Sln1 branch uses a histidine phospho-relay system, which inhibits the kinase Ssk1 in the absence of osmotic stress through the intermediate histidine phosphate transfer protein Ypd1 (Posas et al., 1996). Upon Sln1 activation in response to osmotic stress, Ssk1 inhibition will be relieved resulting in phosphorylation of its downstream MAPKKKs Ssk2 and Ssk22. Like Ste11, these interact and phosphorylate Pbs2, which in turn doubly phosphorylates Hog1, leading to its rapid translocation into the nucleus to launch a transcriptional program. In addition to altered gene expression, particularly induction of Gpd1 (Babazadeh et al., 2014), Hog1-mediated cytoplasmic changes such as the closure of water channels are of great importance to rapidly reestablish osmotic balance (Westfall et al., 2008). Dephosphorylation and inactivation of Hog1 is carried out by an array of phosphatases that includes the tyrosine phosphatases Ptp2 and Ptp3 in the nucleus and cytoplasm respectively, and the Ser/Thr phosphatases Ptc1 and Ptc2/3 (Jacoby et al., 1997; Mapes and Ota, 2004; Murakami et al., 2008; Warmka et al., 2001; Wurgler-Murphy et al., 1997; Young et al., 2002).

The HOG pathway has previously been used to study MAPK cascades as the topology and molecular functions of its components have largely been established (Saito and Posas, 2012). Hog1 activation is traditionally followed by measuring its *in vitro* kinase activity, or phospho-specific antibodies directed against its activating phosphorylation sites or specific Hog1 targets. Moreover, doubly-phosphorylated Hog1 rapidly translocates into the nucleus, which provides a convenient readout to study its activation by quantitative single cell microscopy (Ferrigno et al., 1998). In combination with microfluidic devices allowing precise and rapid control of extracellular osmolarity conditions, such single cell experiments have provided important insight into Hog1 activation kinetics, for example in response to fluctuating inputs (Zi et al., 2010). Finally, mass spectrometry-based studies not only identified numerous downstream Hog1 substrates but also discovered phosphorylation of several HOG pathway components that show drastic changes upon osmostimulation (Kanshin et al., 2015; Vaga et al., 2014) and may thus constitute yet undefined feedback loops (English et al., 2015). Taking advantage of such data sets, previous studies established mechanistic models of the whole HOG pathway (Klipp et al., 2005) or the role of different sub-branches in homeostasis (Schaber et al., 2012), and analyzed the impact of upstream phosphorylation (Hao et al., 2007) or glycerol accumulation (Muzzey et al., 2009) on pathway adaptation. However, experimental data and modeling approaches mechanistically describing Hog1 dual phosphorylation and the relevance of feedback loops for Hog1 activation are still scarce.

In this study, we examined the molecular mechanisms responsible for ultrasensitivity in the HOG pathway. We used an integrative modeling approach taking advantage of various data sources and experimental parameters to compare Hog1 activity and its phosphorylation status under multiple environmental conditions such as varying salt concentrations and salt pulses, and different genetic mutations that specifically perturb Hog1 activation kinetics. Interestingly, our findings support a mixed distributive and processive phosphorylation model as the best fit for the observed experimental behavior, with a critical positive feedback loop targeting the MAPKK Pbs2.

## Results

### Construction of a HOG pathway ODE model comprising putative negative and positive feedback, with different mechanisms of Hog1 activation by double phosphorylation

To understand the molecular mechanism responsible for ultrasensitivity in the HOG pathway, we examined whether the MAP kinase Hog1 is activated by a distributive or processive phosphorylation mechanism. When a basic, three-tiered MAPK module without additional feedback mechanisms is fitted on MAPK activity input-output data ranging from graded to ultrasensitive with unconstrained parameter ranges, we found that parameter sets could be identified that recapitulate the bi-stable behavior irrespective of the phosphorylation mechanism used (Supp Figure S1). We thus constructed an overcomplete model for osmostress-induced Hog1 phosphorylation that allowed evaluating various submodel topologies differing in Hog1 phosphorylation and feedback mechanisms (Figure 1C). A deterministic model based on ordinary differential equations (ODEs) was used due to the relatively high protein concentrations of at least a few hundred molecules per cell, as well as the low variability in the experimental single cell data (Ghaemmaghami et al., 2003; Ho et al., 2018). We applied mass action kinetics to approximate the underlying biochemistry. Taking advantage of previous studies (Zi et al., 2010), we integrated the cell volume module linked to intracellular glycerol concentration, which includes retention and production of glycerol and expression of Hog1-dependent genes (Figure 1C, Supp Figure S2). Downstream effector mechanisms that lead to increased glycerol production and volume adaptation were simplified compared to previous modeling approaches (Klipp et al., 2005) to reduce complexity in areas of the model that are not relevant to explain ultrasensitivity. The model considers both Sho1 and Sln1 branches of the HOG pathway, and the shuttling of Hog1 between a cytosolic and a nuclear compartment. Moreover, it takes into account known and putative feedback regulation, including Hog1-mediated phosphorylation of the upstream components Sln1, Ssk1, Ssk2 and the scaffolding kinase Pbs2 (Sharifian et al., 2015). These reactions result in additional phosphorylated species, which are depicted with new kinetic parameters reflecting their increased or decreased enzymatic activity and/or association rates. Importantly, the model includes both mono- and bi-phosphorylated species of Hog1,and thus allows distinguishing processive and distributive mechanisms for Hog1 activation. In the processive model, the second Hog1 phosphorylation step immediately follows the first without Hog1 disassociating from its scaffold Pbs2, while in the distributive activation model Hog1 disassociates from Pbs2 after the first phosphorylation step and thus needs to rebind to allow formation of the doubly phosphorylated, active species. The distributive mechanism of Hog1 phosphorylation by Pbs2 was implemented by including Hog1-Pbs2 association and dissociation rates. Importantly, the first phosphorylation event always leads to dissociation of the mono-phosphorylated Hog1 species, while in the processive model, the rate constant for the dissociation of monophosphorylated species are set to zero, thus reflecting a case where the Pbs2-Hog1 complex can only dissociate upon double phosphorylation of Hog1. Finally, known phosphatases responsible for Hog1 dephosphorylation (Ptp2, Ptp3, Ptc1, and Ptc2/3) were implemented taking their respective mechanisms into account (Jacoby et al., 1997; Mapes and Ota, 2004; Murakami et al., 2008; Warmka et al., 2001; Wurgler-Murphy et al., 1997; Young et al., 2002). The resulting model thus not only provides a detailed representation of the topology and assembly intermediates upstream of Hog1, but also accounts for different Hog1 activation mechanisms and positive and negative feedback regulation.

### Multimodal data integration and model selection favor distributive over processive mechanism of Hog1 phosphorylation modulated by both positive and negative feedback loops

To infer topology and parameters of the reaction model, we considered experimental data from multiple literature sources, as well as own measurements in wild type and mutant strains exposed to stepwise increase of NaCl of varying concentrations (Figure 1C). The data include population- and single cell measurements directly or indirectly reporting on Hog1 activity at different time points after stimulation (see Supp. Table S1 and S2 for a detailed list and description of considered datasets. For example, mass spectrometry measurements inform about relative changes in single- and double phosphorylated Hog1 within the first 60 seconds of the signaling response as well as at later time points (Kanshin et al., 2015; Vaga et al., 2014). These data were complemented by western blot measurements with antibodies recognizing doubly phosphorylated Hog1 with conditions including strains lacking different upstream components or the Ptp2 and Ptp3 phosphatases, as well as inhibition of Hog1 activity by small molecule inhibitors (English et al., 2015; Jacoby et al., 1997; Macia et al., 2009). Moreover, Hog1 activity correlates with its nuclear translocation, which can be quantified by fluorescence single cell microscopy (Ferrigno et al., 1998). Using a microfluidic platform, we performed extensive Hog1 activity measurements in wild type, *sln1Δ* and *pbs2Δ* cells exposed to various NaCl concentrations and NaCl ramping perturbations. Similar measurements were previously used to assess feedback regulation acting on Ssk2 (Sharifian et al., 2015). Finally, the volume submodel was parameterized with cell area measurements upon various salt treatments of wild type and *pbs2Δ* cells. The latter provides information on Hog1 independent mechanisms that lead to volume adaptation that are also considered in the model (Supp Figure S2).

We used 533 data points across the different conditions and perturbations to estimate the relevant model parameters. We enumerated eight models with distinct topologies varying in Hog1 phosphorylation mechanism and absence or presence of positive and negative feedback loops (Figure 2A). Between 63 and 86 parameters were undetermined in the considered models and thus fitted via likelihood optimization, including the kinetic rate constants of the mass action-based ODEs used to model the general topology and biochemistry of the signaling pathway (see Method section for details).

**Figure 2:**
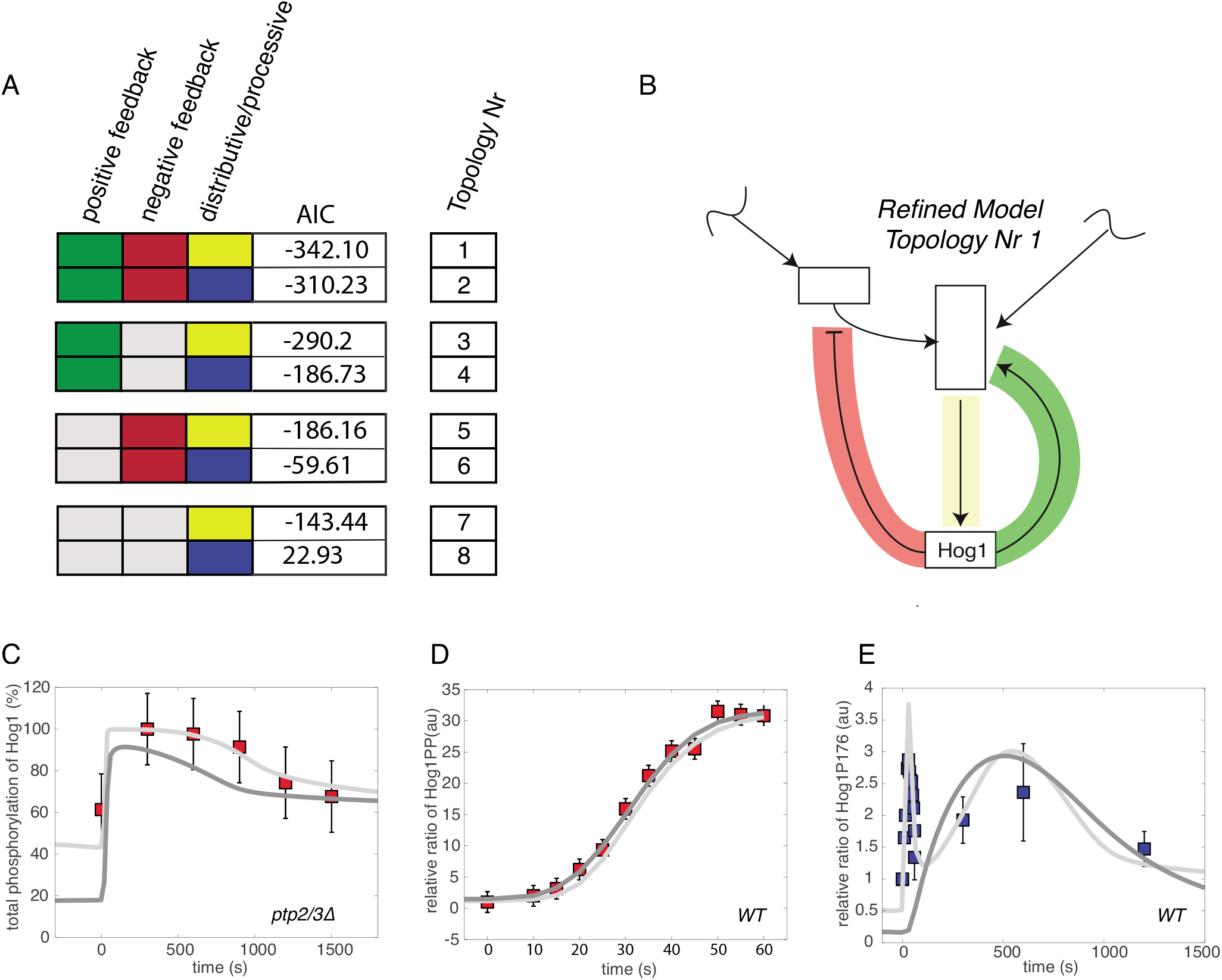
Refined model employs negative and positive feedback and favors a distributive over a processive phosphorylation mechanism. (A) Eight different topologies were enumerated with different combinations of negative feedback (red), positive feedback (green) and distributive (yellow) or processive (blue) Hog1 phosphorylation mechanism. Akaike information Criterion (AIC) of the best model fits are displayed. (B) Abstraction of the model with topology one defined as refined model. Negative feedback on Ssk2 and positive feedback on Pbs2 are indicated in red and green respectively. The distributive phosphorylation mechanism of Hog1 by Pbs2 is visualized in yellow. (C-E) Simulations of selected species of the two best fitting models employing distributive or processive Hog1 phosphorylation (Nr 1 and 2) are shown in response to salt stimulation (0.4M NaCl). Graphs show experimental data points used for parameter optimization (red square), experimental data points used for testing (blue square), simulation results from the best fitting distributive mechanism (light grey) and simulation data from the best fitting processive mechanisms (dark grey). (C) Time course of the total phosphorylation of Hog1 in percent (%) in a Ptp2/3 deleted condition showing simulation and experimental data. (D) Time course of the relative ratio between stimulated and basal levels of dual phosphorylated Hog1 in wild type (WT) cells exposed to 0.4 M NaCl showing simulation and experimental data. (E) Time course of relative ratio between stimulated and basal levels of mono-phosphorylated Hog1-P176 in a WT cells exposed to 0.4 M NaCl showing simulation data and experimental mass spectrometry data that was not used for fitting (Kanshin et al., 2015; Vaga et al., 2014). Note that the processive mechanism does not show the double peak in the data, characteristic of the distributive mechanism.

We first aimed at determining the mechanism of Hog1 activation and evaluating the importance of positive and negative feedback loops. Thus, we performed parameter optimization followed by selection of model variants comprising all combinations of distributive or processive mechanism of Hog1 phosphorylation, and the presence or absence of positive and negative feedback regulation. The Akaike Information Criterion (AIC) was used to compare and rank the results (Figure 2A). Interestingly, comparison of AIC indicated that both positive and negative feedback mechanisms are needed to adequately explain the measurements. In general, an AIC difference between two competing models of greater than ten is considered highly significant (Burnham and Anderson, 2004). Topologies incorporating both feedback mechanisms performed significantly better compared to topologies with only one or no feedback loop, regardless of the Hog1 phosphorylation mechanism. Interestingly, this difference was most pronounced in the case of processive Hog1 activation, where a model without any feedback leads to a break down in model fit, performing significantly worse than its distributive counterparts. Thus, when comparing the two extreme cases of Hog1 phosphorylation mechanisms, the overall best fit was achieved with distributive Hog1 phosphorylation and positive and negative feedback mechanisms, and the AIC difference was significant compared to a processive mechanism. Accordingly, we defined a refined model (Figure 2B). The AIC difference between models utilizing distributive or processive mechanism was most readily apparent in cells deleted for Ptp2 and Ptp3 (Figure 2C). In this case the processive model was unable to recapitulate the full increase in basal signaling as well as the complete activation upon salt stress.

On the other hand, certain finer temporal patterns such as the dynamics of double phosphorylated Hog1 in wild type cells in the first 60 seconds of salt stress, were slightly better approximated by a model employing a processive phosphorylation mechanism (Figure 2D). We interpreted this as a first hint that properties such as the regulation of basal activation levels and maintenance of the full range of activation need a more distributive Hog1 phosphorylation mechanism, whereas certain dynamic properties such as the aforementioned rapid double phosphorylation of Hog1 could more easily be achieved by a processive mechanism.

In an attempt to find additional species that differ significantly between the two best fitting distributive and processive Hog1 activation models and thus might provide some insight into the mechanics underlying their different dynamic properties, we evaluated models for their ability to explain experimental measurements that were not part of the initial simulations. We found some phospho-species that show very distinct behavior. In particular, simulations of mono-phosphorylated Hog1 dynamics revealed significant qualitative differences upon simulation with different phosphorylation mechanisms (Figure 2E). Simulations with a distributive phosphorylation mechanism resulted in a temporal profile marked by a double peak, with a first activity peak during the initial minutes and a second, lower increase towards the end of the response. In contrast the processive model simulation displayed monophosphorylated Hog1 dynamics with a single peak reminiscent of the behavior of doubly phosphorylated Hog1. Since the mono-phosphorylation data were not used for the parameter fitting process, these Hog1 activation dynamics are likely an inherent property of the system and not the result of overfitting.

### A mixed phosphorylation mechanism best explains Hog1 activation kinetics by integrating favorable dynamic properties of both distributive and processive mechanisms

While distributive Hog1 activation leads to a significantly better goodness-of-fit, the processive mechanism better recapitulates some data such as speed and degree of double phosphorylated Hog1 accumulation. Thus, we next evaluated a mixed phosphorylation mechanism that incorporates characteristics from both the processive and distributive mechanism. This mixed mechanism allows for both processive double phosphorylation and distributive dissociation of mono-phosphorylated species (Figure 3A). Thus, monophosphorylated Hog1 has a certain propensity, defined by the kinetic rate constant, to remain Pbs2 bound and immediately undergo a second phosphorylation step, in which case we observe processive characteristics. Alternatively, mono-phosphorylated Hog1 can dissociate from Pbs2 resembling a distributive mechanism. To assess whether such a mixed model would indeed recapitulate above finer temporal patterns of Hog1 activation, we repeated the parameter optimization procedure by including mass spectrometry data sets of monophosphorylated Hog1 the results of which we defined as best fitting models (Kanshin et al., 2015; Vaga et al., 2014) (Supp Figure S3). Interestingly, AIC measurements revealed that the optimized model employing a mixed phosphorylation mechanism showed significantly better goodness of fit compared to the either distributive or processive variants, with all models incorporating positive and negative feedback (Figure 3B).

**Figure 3:**
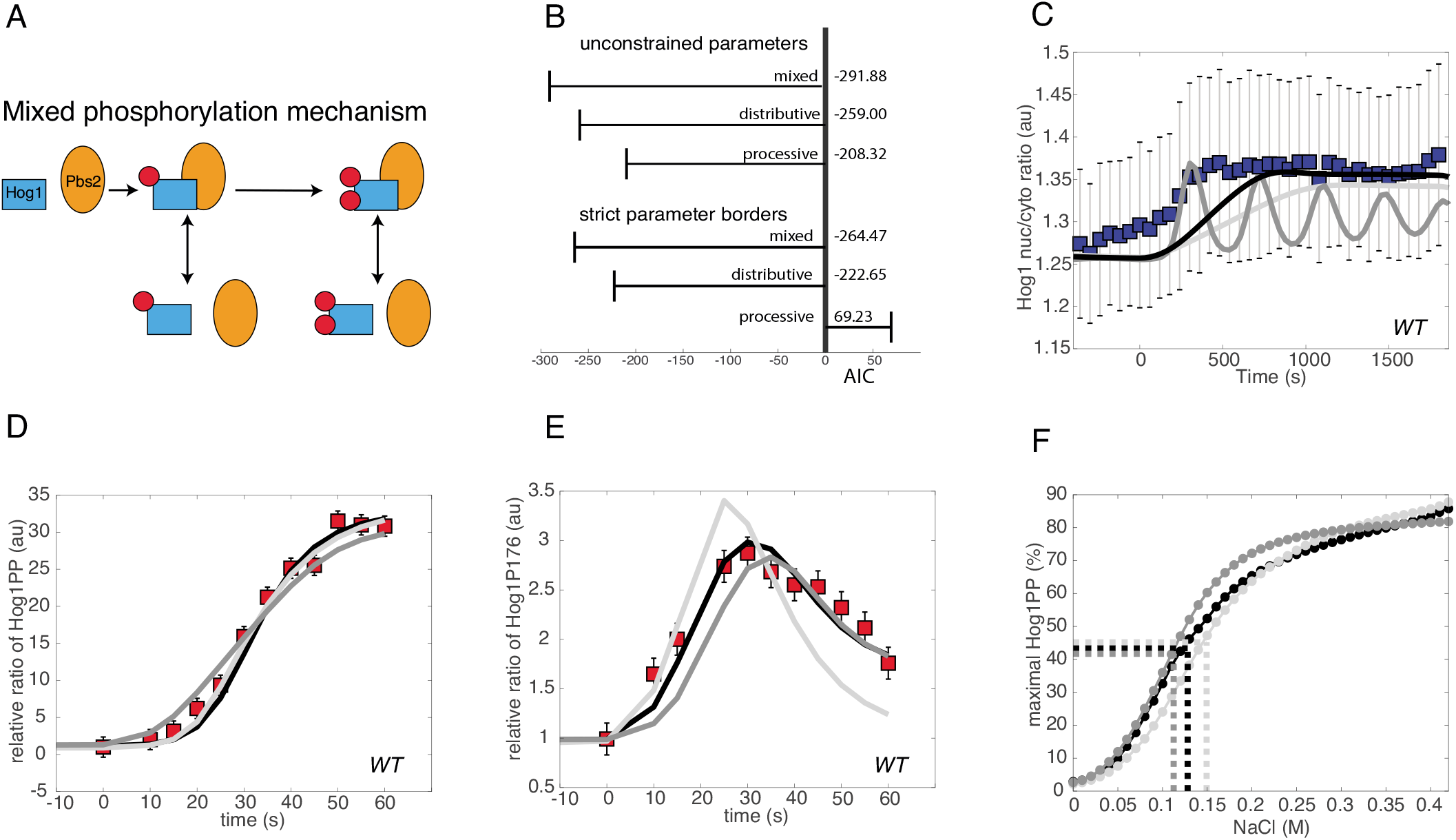
Temporal dynamics of monophosphorylated Hog1 supports models with partially distributive phosphorylation. (A) A mixed phosphorylation mechanism generates a phosphorylation reaction with both distributive and processive character. (B) Akaike information criterion (AIC) of the best fits after the optimization and fitting with unconstrained parameter values or stricter boundaries that approximate physiological conditions (see Supp Fig S4). Note, that in contrast to the refined model of Figure 2B, data from mono-phosphorylated Hog1 species was used in the fitting and the resulting fit thus the best fitting model on all data. (C-E) Simulations are shown of selected Hog1 species from the three best fitting models in which all models incorporate positive and negative feedback in wild type cells (WT). Graphs show experimental data points used for parameter optimization (red square) and testing (blue square), as well as simulation results from the best fitting mixed (black line), processive (dark grey line) or distributive (light grey line) Hog1 phosphorylation mechanism in response to salt stimulation. (C) Time course of Hog1 nuclear to cytosolic ratio in cells exposed to linear ramping up to 0.2M NaCl showing simulation and experimental data. Note that the experimental data (blue squares) was not used in the fitting procedure. (D – E) Time course following salt stimulation (0.4M NaCl) of mono-phosphorylated Hog1-P176 (D) and dual phosphorylated Hog1-PP (E) expressed as a relative ratio between stimulated and basal levels showing simulation and experimental data. (F) Fit of a Hill function to the simulated input-output curves of the best fitting mixed (black line), processive (dark grey line) or distributive (light grey line) Hog1 phosphorylation mechanism with a Hill coefficient of 2.39, 3.03 and 2.77, respectively. EC50 values are indicated by a dashed line of respective colors.

We used the best fitting distributive, processive or mixed models with unconstrained parameters to predict Hog1 activation upon a linear ramping increase of NaCl concentration over a prolonged period of time, comparing simulation output to an independent data set not used for parameter optimization (Figure 3C). Monitoring Hog1 nuclear translocation dynamics to approximate Hog1 activation, the processive mechanism leads to premature Hog1 translocation and displays an inappropriate oscillating behavior, while the distributive mechanism results in a delay of Hog1 translocation. Only the mixed model was able to finetune the Hog1 nuclear translocation kinetics with an appropriately rapid yet stable response.

Indeed, the processive mechanism produces premature accumulation of activated Hog1 manifested by the appearance of doubly phosphorylated Hog1 in the first 60 seconds after salt addition (Figure 3D). We assume that this behavior addresses the need to compensate direct mono-phosphorylation by employing faster accumulation of double phosphorylated Hog1 that can then undergo dephosphorylation to generate the pool of mono-phosphorylated species. Case in point, the simulated accumulation of monophosphorylated Hog1 by the processive model was slower and not as high as expected from experimental data (Figure 3E). On the other hand, the distributive model resulted in faster accumulation of mono-phosphorylated species than measured experimentally and its decay set in earlier than expected (Figure 3E). However, the mixed phosphorylation mechanism shows a better fit and is situated between the two extreme cases, and thus as predicted compensates for both premature accumulation and decay observed with distributive Hog1 activation and the delay of the processive mechanism (Figure 3D, E). Upon fitting Hill functions to our simulated input-output curves, we observed that increase in processivity led to lower EC50 values, with the mixed mechanism achieving a lower value than a distributive and a processive mechanism. We also quantified ultrasensitivity using the Hill coefficient (Figure 3F). Interestingly, the processive mechanism resulted in the highest Hill coefficient, while as expected, the mixed mechanism showed lower ultrasensitivity than the distributive mechanism.

We further considered physiological parameter boundaries more closely reflecting known parameter values of general yeast kinases and phosphatases. Within these boundaries the mixed phosphorylation mechanism resulted in an even more significant difference in AIC compared with the extreme mechanisms (Figure 3B). For example, with physiological parameter boundaries the processive mechanism was unable to recapitulate the double peak behavior of mono-phosphorylated Hog1 and experimental data specific to the Sho1 subbranch of the pathway (Supp Figure S4).

### Hog1-dependent positive feedback increases processivity of the Hog1 phosphorylation reaction

Next, we investigated why the mixed Hog1 activation model best fits the observed measurements. Our model simulations confirmed that positive feedback is necessary to achieve the best goodness of fit (Figure 2A). This becomes most readily apparent with data quantifying Hog1 double phosphorylation when its kinase activity is inhibited by small molecules (Supp Figure S3, Cond 16) (English et al., 2015; Macia et al., 2009). Positive feedback is necessary to recapitulate both the slow Hog1 phosphorylation upon kinase inhibition as well as the rapid and full activation of Hog1 in wild type cells.

To explore targets and mechanisms underlying this positive feedback, we analyzed mass spectrometry measurements. Indeed, these measurements revealed that eight out of eleven components involved in the Hog1 pathway undergo phosphorylation upon salt exposure (Vaga et al., 2014). We focused on putative feedback loops that phosphorylate targets upstream of Hog1. In the Sln1 sub-branch, the data suggests that Ssk1 and Ssk2 are potential Hog1 substrates, while in the Sho1 sub-branch Hog1 phosphorylates Ste50 and Ste20. However, phosphorylation of Ssk2 (Sharifian et al., 2015) and Ste50 (Yamamoto et al., 2010) interfere with Hog1 activation, making them unlikely physiological candidates. Moreover, positive feedback must target both sub-branches simultaneously to explain the experimental data. Therefore, Hog1-dependent phosphorylation of Pbs2 would fulfill this requirement, as Pbs2 integrates the information of both sub-branches (Figure 4A). Alternatively, we considered Ste20 and either Sln1 or Ssk1 as potential targets of positive feedback. Interestingly, a mixed model that incorporates positive feedback of Hog1 on Pbs2 showed significantly better results than topologies in which Hog1 targets Sln1 and Ste20 or Ssk1 and Ste20 (Figure 4B). The changes introduced by the positive feedback mainly lead to increased processivity, meaning enhanced speed of the second Hog1 phosphorylation reaction with only minor effects on the first phosphorylation reaction. Processivity in the mixed Hog1 activation model is formally defined by the ratio between the rate of the second phosphorylation step and the rate of dissociation of mono-phosphorylated Hog1 bound to activated Pbs2 (Figure 4C). To assess processivity over the course of the response we introduced a processivity score (see Method section), which quantifies the rate of an immediate second phosphorylation step compared to dissociation of the Hog1-Pbs2 complex. Interestingly, the simulations showed that in the best fitting model with positive feedback on Pbs2, this score is low before and after the response to salt stimulation and the Hog1 phosphorylation mechanism displays a behavior mimicking a distributive mechanism (Figure 4D). The onset of positive feedback activity during the response leads to a significant increase in the score, meaning the reaction becomes highly processive. In comparison, the best fitting result of the model targeting Sln1 does not result in significant changes of processivity. The Ssk1 model displayed increased processivity along the course of the reaction but the basal level of processivity was orders of magnitude higher than the better fitting model targeting Pbs2. Taken together, these results indicate that mixed Hog1 phosphorylation enables a switch between a distributive and processive activation mechanism. This switch is initiated by positive feedback, most likely regulating Pbs2.

**Figure 4:**
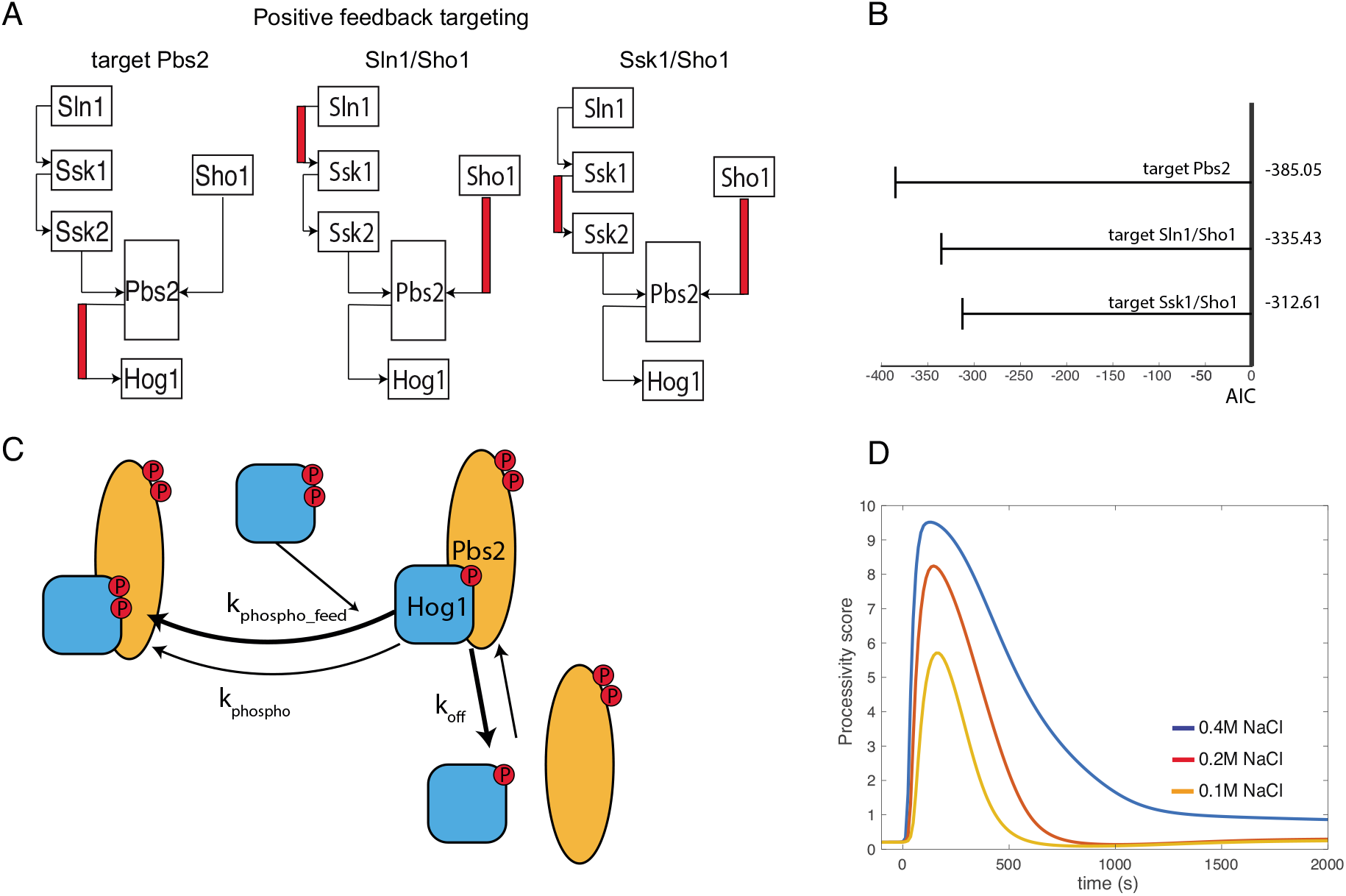
Positive feedback to Pbs2 gives best goodness of fit and results in increase of processivity. (A) Schematic of three models with potential targets for positive feedback upstream of Hog1, showing the positive feedback highlighted in red for models where activated Hog1 phosphorylates either Pbs2 (left), Sln1 and Sho1 (middle), or Ssk1 or Sho1 (right). (B) Akaike information criterion (AIC) was used to order optimized models with the indicated positive feedback targets, in which all models include negative feedback and a mixed Hog1 phosphorylation mechanism. Positive feedback targeting Pbs2 achieves best AIC among other putative targets. (C) The phosphorylation of Pbs2 by activated Hog1 introduces a second, faster rate constant k_phospho_feed_ for the activation of Hog1. The rate constant of Hog1-Pbs2 dissociation remains unaffected. (D) Time course upon salt stimulation showing a processivity score that quantifies the ratio between the rate of the second Hog1 phosphorylation step and its dissociation from Pbs2 (see Methods). A low processivity score indicates distributive-like phosphorylation, while a high processivity score indicates a processive-like mechanism. Note that before starting the reaction the processivity score is low, but the onset of Hog1 signaling introduces positive feedback (t=0), which increases the score until the reaction behaves like a processive mechanism.

### Mixed mechanism conveys robustness to protein level fluctuations

We next evaluated robustness for the mixed Hog1 activation mechanism, specifically determining to what degree a similar output can be generated across a spectrum of external or internal perturbations. Robustness is of particular importance in the case of Hog1 phosphorylation since its hyperactivation leads to severe growth defects, observed for example upon overexpression of Pbs2 (Krantz et al., 2009). Indeed, *in silico* overexpression of Pbs2 revealed that Hog1 was almost fully doubly phosphorylated and thus hyperactivated in the best fitting distributive and mixed models (Supp Figure S5A). In contrast, overexpression of Pbs2 in the processive model actually prevented double phosphorylation of Hog1 (Supp Figure S5B). To corroborate these findings, we randomly varied the concentration of Hog1 and protein species upstream of Hog1, as well as varying the Hog1 phosphatases in a more narrow range of half to twice their original concentration (Figure 5A). We repeated these simulations over 500 times and examined the maximal activation in response to various concentrations of salt stimulation of the best fitting distributive, processive and mixed models (Figure 5B-D). With no perturbation or low NaCl concentrations (0 M, 0.1 M), the processive mechanism resulted in slightly bimodal distributions with 2.4 times the number of perturbations resulting in complete activation (> 70% Hog1 double phosphorylation) than the mixed mechanism and the majority showing no activation. The distributive mechanism led to an accumulation of activation at an intermediate level (20%), while the mixed model distributed more around the unperturbed activation level (Figure 5B, C). At higher levels of NaCl input (0.3M), the processive mechanism failed to reach the maximal level of activation and displayed a disproportionate amount of simulations that led to only very minimal activation. The distributive mechanism on the other hand, shows a more uniform distribution between 30 and 90 % Hog1 activation. Strikingly however, only the mixed mechanism predicted low basal levels of Hog1 activity, with reliable rapid and high activation upon exposure to the different NaCl conditions (Figure 5D). Moreover, multiple linear regression suggests that increasing Ssk1 and to a lesser degree Ssk2 concentration most significantly contributed to erroneous Hog1 activation at lower (0, 0.1M) salt concentrations. With higher salt input (0.3M) changes in Pbs2 concentration become more important to explain the changes in output (Figure 5E). We thus conclude that a mixed Hog1 activation model regulated by a positive feedback loop fits the complete, consolidated data significantly better than either purely distributive or processive mechanisms. Moreover, the mixed Hog1 activation displays favorable systems behavior and leads to increased robustness in the inputoutput relations.

**Figure 5:**
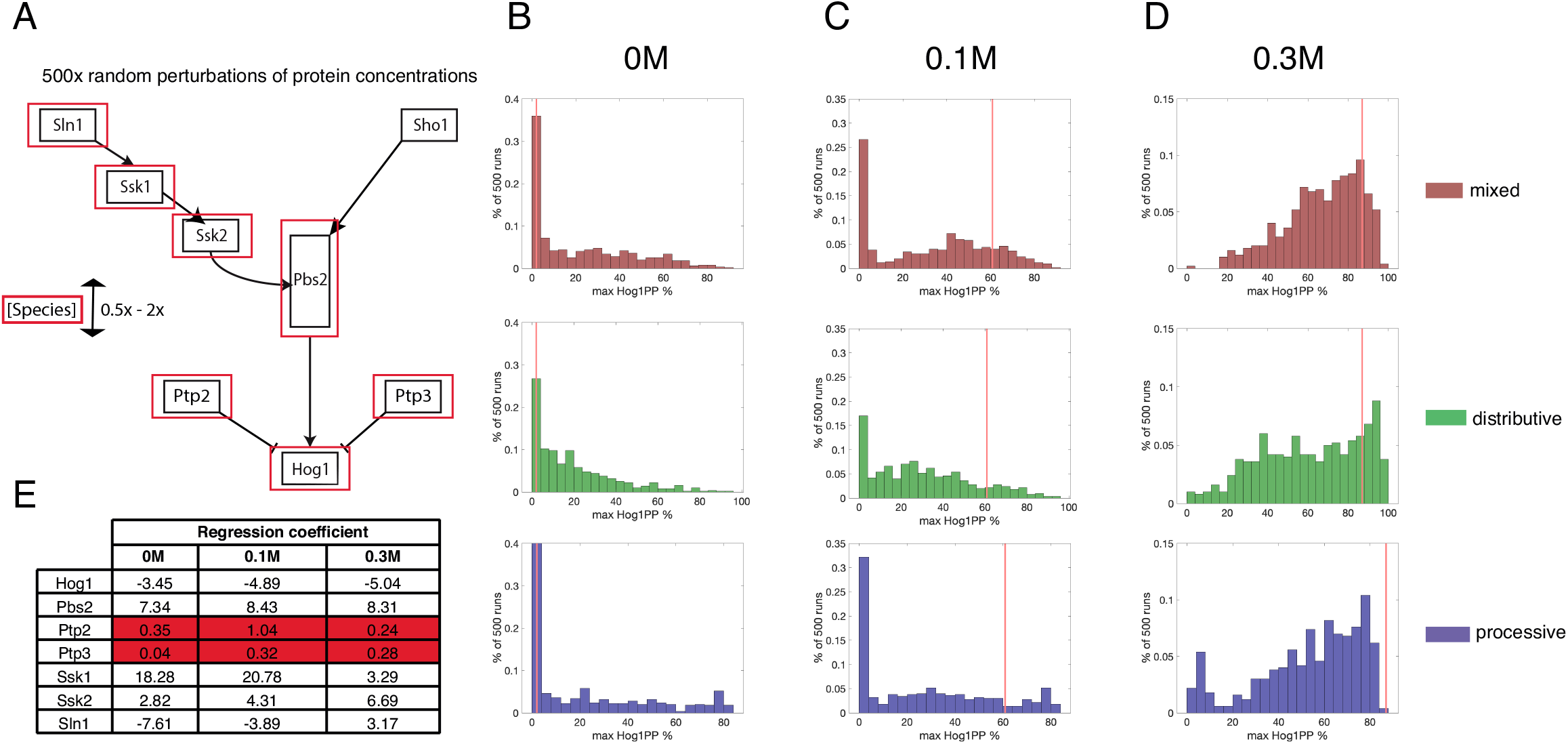
Mixed mechanism with positive feedback is more robust to protein concentration perturbations. (A) Schematic of the Hog1 pathway indicating selected species (red box) randomly varied in concentration by 0.5 to two times their physiological concentration, generating 500 new concentration vectors. (B – D) The best fitting models of mixed (red), distributive (green) or processive (blue) phosphorylation mechanism incorporating the concentration vectors generated as in (A) were used to simulate the maximal activation of Hog1 under different salt concentrations. The red line indicates the maximal activation of the best fitting unperturbed model. (B) With no salt perturbation, all models show a majority of runs with no Hog1 activation as expected. The processive model shows a portion of runs that result in full activation. The distributive model accumulates activation of around 20 percent, while the mixed model shows less accumulation of fully or partially activated Hog1. (C) At low salt stimulation (0.1M NaCl) all models show a majority of simulations that tend to non-activation. The processive mechanism shows a sizeable portion of erroneously fully activated runs, while runs of the distributive mechanism tend to accumulate at 20 percent. The mixed model clusters better around the correct percentage of activated Hog1. (D) At higher salt stimulation (0.3M NaCl), all models show a majority of runs with full activation. The processive mechanism displays a bimodal distribution with a sizeable number of runs that are not activated. The distributive mechanism results in a broader, nearly uniform distribution between 30 and 90 percent. The mixed mechanism results in runs distributed around the correct full activation with fewer runs showing non-activation. (E) Normalized regression coefficients for each of the changed species as a result of multiple linear regression. Higher values indicate higher contribution to explaining the variance of the data. Non-significant coefficients (p-value < 0.05) are highlighted in red.

## Discussion

Effective processing of external information via subsequent intracellular response is of vital importance for every cell. From a systems-engineering perspective it is desirable for this process to be robust towards external and internal state fluctuations. Similarly, the actual input threshold to induce a cellular response should be carefully chosen to avoid unnecessary energy expenditure. It can be argued that this is specifically true for stress response pathways that need to react rapidly and often have dramatic effects on the cellular state. In the *S. cerevisiae* HOG pathway, Hog1 activity not only triggers rapid accumulation of intracellular glycerol but also inhibits other signaling responses such as cell cycle progression and mating (Escote et al., 2004; O’Rourke and Herskowitz, 1998). Thus, while Hog1 activity must be kept low in the absence of osmolarity stress, fast Hog1 activation is required to prevent cell lysis. However, Hog1 activity also needs to remain readily reversible to ensure that cell growth can resume after the pressure difference is balanced. Control over the system can be exerted by feedback loops that are able to tune the signaling response. Moreover, like other MAP kinases, Hog1 is activated by dual phosphorylation and thus the choice and kinetics of this phosphorylation mechanism can greatly affect signaling dynamics. A multitude of previous, mostly theoretical, work has focused on the extreme cases of distributive or processive mechanisms. Interestingly however, our combined experimental and modeling analysis revealed that a mixed control mechanism regulated by a positive feedback loop best explains the observed phosphorylation and signaling dynamics. Available data, however, cannot exclude that lower diffusion rates caused by cell shrinkage may contribute to increased processivity (Babazadeh et al., 2013; Miermont et al., 2013). Indeed, it has been argued that molecular crowding might affect processivity in HeLa cells (Aoki et al., 2011). However, since the possibility of tuning MAPK activation results in specific and emergent systems output, regulated switching between a distributive and processive phosphorylation mechanism is best explained by positive feedback.

### Positive feedback may be needed to buffer noise and crosstalk to allow high basal signaling

In mammalian cells positive feedback loops have been recognized and described in multiple pathways. In the ERK pathway for example, positive feedback loops have been identified that trigger an oscillatory behavior in combination with negative feedback (Kochanczyk et al., 2017). The distinction between prolonged or oscillatory response and its exact frequency is crucially important to determine the cellular output (Ryu et al., 2016). In contrast, in the yeast *S. cerevisiae* stress response pathways, no such oscillating signaling dynamics are known. The ability to buffer noise while retaining the sensitivity of the reaction could provide an evolutionary benefit (Hornung and Barkai, 2008). Indeed, there is considerable basal signaling from the Sln1 sub-branch of the HOG pathway (Macia et al., 2009). A positive feedback loop might thus be needed to set a threshold of Hog1 phosphorylation that is not crossed without increased activation of the pathway by exposure to hyperosmolarity. Similarly, positive feedback could play an important role to prevent erroneous activation of Hog1 by crosstalk from other pathways such as the pheromone response pathway (O’Rourke and Herskowitz, 1998).

However, no positive feedback mechanisms regulating the HOG pathway have been described at present. According to our models the most likely target for such a positive feedback regulation is Pbs2. Indeed, Pbs2 is phosphorylated on S248 in response to high osmolarity (Kanshin et al., 2015; Vaga et al., 2014) and this serine is followed by a proline residue and thus conforms to the minimal MAPK consensus motif (S/TP). However, inhibition of Hog1 kinase activity does not seem to attenuate S248 phosphorylation (Romanov et al., 2017). Interestingly, recent experiments confirm osmostress-mediated enhancement of the reaction between Pbs2 and Hog1 (Tatebayashi et al., 2020), and suggest that a new downstream osmosensor could fulfill the role of positive feedback. Thus, further work will be required to pinpoint positive feedback mechanisms regulating HOG signaling.

### Mixed mechanism may finetune sensitivity and work in tandem with positive feedback to guarantee a well-defined threshold and robustness

In addition to positive feedback, our best fitting model also displays both distributive and processive qualities. Previous theoretical analyses of a simple three-tiered MAPK cascade predicted that such a mixed mechanism may enhance the tunability of the response (Sun et al., 2014). In particular, increased processivity can enhance the sensitivity to the external signal and lower ultrasensitivity. Indeed, increasing processivity diminished EC50 values, with the mixed mechanism achieving a lower value than distributive and processive mechanisms (Figure 3F). Moreover, measurements of the Hill coefficient confirmed that the mixed mechanism showed lower ultrasensitivity than distributive Hog1 activation. This is probably due to the processive mechanism plateauing earlier with a lower maximal activation than the two other mechanisms. This would indicate that while a processive mechanism can guarantee ultrasensitivity, the range of dynamics is lower than what a distributive mechanism can achieve. Despite this effect being significant, its physiological relevance under normal conditions appears small. However, under conditions where the difference in the inputoutput curves becomes more pronounced, this mechanism may become important to ensure cell survival.

A bigger impact of the different phosphorylation mechanism was observed when evaluating robustness of the pathway output. The mixed mechanism was more robust towards perturbation of protein concentrations, and also less prone to exaggerated activation at lower salt concentrations. Multiple linear regression suggests that in particular increasing Ssk1 or Ssk2 levels contribute to maximal Hog1 activation at basal input levels (Figure 5E). Available experimental data confirms that the Sln1 sub-branch displays high basal signaling (Macia et al., 2009). We propose that the distributive nature of Hog1 phosphorylation with or without low salt stress serves as an additional checkpoint, where mono-phosphorylated Hog1 species are dephosphorylated faster than the second phosphorylation step and thus full Hog1 activation can occur. However, after a certain input threshold has been crossed at higher salt concentrations, the positive feedback increases the processivity of the reaction, reliably promoting maximal Hog1 activation. In comparison, the purely distributive mechanism resulted in 5.5% lower activation on average at the highest salt concentration.

Our findings on the processivity of Hog1 phosphorylation may have important implications to explain dynamic properties of MAPK pathways in mammalian systems. For example, the mammalian Hog1 homolog p38 was recently shown to be phosphorylated in a semi-processive manner (Wang et al., 2019), which may be best explained by a mixed activation model. Likewise, mammalian ERK shows context-specific differences in its ratio of single and double phosphorylated species (Iwamoto et al., 2016).

### The tunable phosphorylation mechanism with positive feedback: a general mechanism for multisite phosphorylation?

Our results imply that the switch like behavior and ultrasensitive response of MAPK modules is a function of the whole cascade. Interestingly, other switch-like transitions such as cell cycle progression or cell differentiation use an increasing number of phospho-sites of effector proteins as a strategy to alleviate the need for a cascade. For example, degradation of the *S. cerevisiae* cyclin-dependent kinase inhibitor Sic1 requires phosphorylation of at least six residues to allow S phase entry (Nash et al., 2001). It has been argued that a distributive mechanism to phosphorylate these sites results in a highly ultrasensitive, switch-like response. Sic1 is phosphorylated by two distinct kinase complexes, Cln2-Cdk1 and Clb5-Cdk1, which bind and first phosphorylate priming phospho-sites on Sic1, which in turn triggers processive phosphorylation of the entire phospho-degron (Koivomagi et al., 2011). This systems behavior could thus be viewed as an extreme case of a mixed phosphorylation mechanism, switching from a distributive to a processive mode of activation. Theoretical analyses predicted that increasing the number of phosphorylation sites sharpens the threshold, but might only allow for a graded increase beyond the threshold (Gunawardena, 2005). However, a theoretical framework with a mixed mechanisms confirms its potential for additional behavior already in a system with three phosphorylation sites (Suwanmajo and Krishnan, 2015). Indeed, experimental evidence for a mixed mechanism termed semi-processive phosphorylation was described for the multisite phosphorylation of Pho4 (Jeffery et al., 2001). It is thus tempting to speculate that the here described tunable phosphorylation mechanism in combination with regulatory feedback could be utilized not only to activate MAPK’s but may more generally apply to many multisite phosphorylation systems, as the resulting properties are robust and able to generate ultrasensitive signaling dynamics such as efficient switching and threshold adaption.

## Supporting information

Supp model files

## Acknowledgements

We thank S. Pelet (University of Lausanne) for the SKAR reporter and B. Hegemann and F. van Drogen for help with strain construction. Numerical simulations were performed on the ETH Euler cluster. We are grateful to A. Smith for critical reading of the manuscript and members of the Claassen- and Peter laboratories for helpful discussions. Research in the Peter lab was funded by the Swiss National Science Foundation, the European Research Council (ERC), ETH Zurich and the Global Research Laboratory (NRF-2015K1A1A2033054) through the National Research Foundation of Korea (NRF). Research in the Claassen lab was supported by internal funds of ETH Zurich.

## Author Contributions

MM developed the modeling framework and conducted the computational analysis; MM and SSL conducted the experiments; all authors participated in study design and wrote the paper.

## Methods

### Yeast strains, microscopy and microfluidics

Yeast strains and plasmids are listed in Supplementary Tables S1 and S2. yMU49, which provides the nuclear marker HTA2-CFP, was used as a starting strain. HOG1-YFP was amplified by PCR from yMU19, and a simple transformation protocol using lithium acetate, polyethylene glycol (PEG) and heat shock induced transformation was utilized to create yMM001. The SKARS sensor (Durandau et al., 2015) or dPSTR (Aymoz et al., 2016) were transformed into yMM001 after cutting of plasmids pED45 or pDA183 with corresponding restriction enzymes to create yMM003 or yMM004 respectively. The Pbs2 deleted strain was created by transformation of the PCR amplification product of the NAT cassette from pSP135 with primers containing sequences 1000 bp up- or downstream of the gene of interest. Successful plasmid cut and PCR product length was confirmed by gel electrophoresis.

For live cell imaging experiments, yeast cells were grown in Synthetic Defined (SD) media with 2% glucose. Exponentially-growing cells were transferred to a microfluidic device (Y04C, Merck Millipore) and live cell imaging carried out at 30°C using a fully-automated inverted epi-fluorescence microscope (Ti-Eclipse, Nikon) in an incubation chamber. Osmotic stress was induced with the pressure controller (ONIX, Merck Millipore) by exchanging the medium with media containing NaCl at the specified concentrations. The images were taken with a high numerical aperture oil immersion objective lens (CFI Plan Apo 60X, Nikon; N.A. =1.4). Image acquisition was controlled using micro-manager. Each frame was imaged with relevant fluorescent set-up (CFP, YFP, mCherry and Cy5 fluorescent filters with LED illumination). Cell segmentation, tracking and feature extraction was done using the MATLAB^®^-based YeastQuant software (Pelet et al., 2012), using Alexa 680 fluorescent dye for cell segmentation (Pelet et al., 2012). The CFP channel was used to define cytosolic and nuclear regions based on HTA2-CFP images, by defining a certain intensity threshold. Individual cells during time-lapse imaging were followed by tracking the nucleus. The cytosol and nucleus of individual cells was segmented and various properties including cell area and average intensity of fluorescent signals in the segmented objects were quantified.

### Data sources and incorporation into model definition

The experimental data was collected from multiple sources. When not otherwise noted the BY4741 yeast strain exposed to 0.4 M NaCl stress was used. Importantly mass spectrometry measurements analyzing long term (Vaga et al., 2014) and very short term (Kanshin et al., 2015) changes in phospho-site abundance were included. Since the relative phospho-site change at the timepoint 60 seconds after NaCl addition varies slightly between these two studies, the mean of both values was taken for this time point.

To determine the degree of Hog1 phosphorylation, the maximal values determined by western blot measurements were used (English et al., 2015). Data for Hog1 phosphorylation upon inhibition of Hog1 with or without addition of salt was taken from the same experimental set. Even though the authors used a different strain with a different perturbation agent (KCl instead of NaCl) the qualitative dynamics of the Hog1 response were essentially identical (Muzzey et al., 2009). Moreover, the switch-like response resulting in full Hog1 phosphorylation for all salt concentrations above a certain, low threshold justified inclusion of these data.

Additional western blot measurements were utilized for different phosphatase mutants. Compared to mass spectrometry measurements, quantification of western blots is complex and these results often reveal more qualitative than quantitative results (Taylor et al., 2013). For example, different antibodies are known to have different binding properties (Tinti et al., 2012). As a consequence, measurements of Hog1 phosphorylation in Ptp2/Ptp3 mutants in the literature revealed quite varying results (English et al., 2015; Jacoby et al., 1997; Mattison and Ota, 2000; Murakami et al., 2008; Wurgler-Murphy et al., 1997). However, despite these differences, the data largely agree that deletion of Ptp2 results in increased basal levels and prolonged Hog1 phosphorylation, while deletion of Ptp3 shows only minute differences compared to wild type controls. To account for this variability, we utilized the data of Jacoby *et al.* (1997), but allowed the parameters for the error model of these measurements to be sufficiently big and independent from other western blot data sets. Furthermore, as the used antibody used detects phosphorylated tyrosine, the corresponding output species of the model was set to be Hog1 doubly phosphorylated or mono-phosphorylated at tyrosine 176.

Microscopy measurements of Hog1 relocation to the nucleus report on Hog1 activity, and such data was used to quantify the response of single branch and phospho-site mutants. Similarly, microscopy-based measurements of cell size were used to fit the volume module. Information on Hog1 activation at 0.2 M NaCl stimulation with varying frequency (2, 4, 8, 16 min) were extracted from (Mettetal et al., 2008). Volume and Hog1 relocation measurements in cells deleted for *SLN1* or *SHO1* were taken from (Granados et al., 2017). Although the authors used sorbitol instead of NaCl for their experiments, own measurements confirmed that the response of cells to 0.6 M sorbitol is quantitatively and qualitatively nearly identical to perturbations with 0.4 M NaCl. To assess negative feedback of Hog1 on Ssk2, we used published Hog1 relocation measurements (Sharifian et al., 2015). Due to the fact that non-phosphorylatable Ssk2 mutant exhibit reduced cell shrinkage and thus a lower maximal Hog1 nuclear to cytosolic ratio, only data from later time points were included, which correct for the increased time it takes for the Hog1 ratio to reach basal levels. Data for Gpd1 expression changes were selected from (Rep et al., 1999), and information about Ptp2 and Ptp3 mRNA levels were extracted from (Wurgler-Murphy et al., 1997). Western blot measurements of total Hog1 phosphorylation upon inhibition with small-molecule inhibitors in wild type and *ssk2D* strains were taken from (Macia et al., 2009) for basal activity and (English et al., 2015) for activity upon increase of osmolarity. As most antibodies used to measure Hog1 phosphorylation bind to both the doubly and mono-phosphorylated species (English et al., 2015), these western blot measurements were considered as total amount of phosphorylated Hog1 irrespective of phosphorylation grade.

The observables in these data were incorporated as model variables by defining observables that correspond to the experimentally measured quantity. We defined the relative ratio between the current absolute number of a protein species and its absolute number at the very start, the phosphorylated percentage as the absolute number of phosphorylated protein divided by the total number of the protein, and the concentration ratio of Hog1 in the nucleus compared to the cytosol, for the mass spectrometry, western blot, and fluorescence ratio data sets respectively. Further details on how the observables and their corresponding data set were encoded in the model can be gathered from Supp Table S5: Observables.

### Modeling

All modeling was performed using the Data2Dynamics modeling environment (Raue et al., 2015) and computation was carried out on the Euler cluster provided by ETH Zurich. Parameter optimization was carried out by multistart followed by a deterministic trust region algorithm. The goodness of fit was evaluated by Goodness-of-fit = −2 * *log*(Likelihood). For parameter optimization, parameters were considered on a log-scale. In a first step, 10000 starting parameter vectors were created. Sampling was done uniformly between the lower and upper boundaries of each parameter. Lower and upper boundaries were first set to be minus three to three, respectively, spanning six orders of magnitude. After successful parameter estimation (after either 400 iterations or if the value of the objective function changes by less than 1e-6) the 100 best fitting parameter vectors were taken and their correlation coefficient determined. 10000 starting vectors for the next optimization run were generated by sampling from a multivariate normal distribution whose parameters were determined by the 100 best fitting parameter vectors of the previous run. This procedure was repeated until the best goodness of fit of the latest run turned out as good or worse than the run before.

To minimize the risk of the optimization procedure redundantly reproducing solution clusters with very similar values, we included additional steps, such as optimization of crucial model topologies being redone up to eight times with different starting parameters, and ensuring that the relative ordering of the fits was consistent over various conditions, such as leaving out of data points, or changing of parameter boundaries. In general, parameter identifiability was not a major concern, as some model topologies were not able to recapitulate the data and we were primarily interested in the relative quality difference of each topology prediction. The size of the model and its complicated nature also impeded identifiability analysis by profile likelihoods. Instead we simulated species concentration from a model with known parameters. From these simulations we sampled data points equivalent in number and time points to the experimental data set, and performed the parameter optimization procedure described above. All but twelve of the 82 newly fitted parameters were within one order of magnitude from the parameters used to generate the data, with a median of 0.11.

In all our models, certain parameters of the Sho1 sub-branch (Sho1 binding to Ste11, phosphorylation of Pbs2 by Sho1 and dissociation of activated Pbs2 from Sho1) assumed values much higher than expected (up to 10^6) after parameter optimization. We speculate that this reflects a stable complex that forms at the cell membrane which incorporates all involved species (Sho1, Ste11, Ste50, Ste20, the scaffolding protein Ahk1, and more) and brings them into close proximity so that reactions occur faster than expected by simple diffusion (Nishimura et al., 2016; Takayama et al., 2019; Truckses et al., 2006). As this was observed with all model topologies, we feel confident to compare the different topologies relative to one another even if the models do no fully capture the Sho1 sub-branch. Similarly, we did not incorporate the latest information that describes how Ste11 only phosphorylates one phospho site on Pbs2 (Tatebayashi et al., 2020).

Comparison of different models was done using the Akaike Information Criterion.

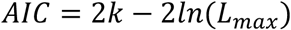

With “k” being the number of fitted parameters and *L_max_* the maximal likelihood.

Detailed information on the best fitting mixed model can be found in Supp table S6-S9.

### Processivity score

We introduced a score (1) to quantify the change in processivity of the Hog1-Pbs2 reaction over the course of the response.

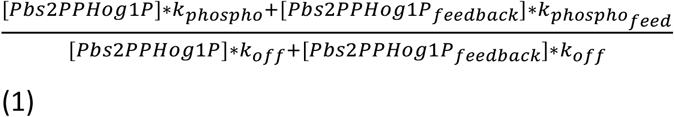

It is defined as the probability of a monophosphorylated Hog1 bound to activated Pbs2 undergoing a second phosphorylation step against the probability of the proteins dissociating. Feedback on Pbs2 is reflected in a higher rate of phosphorylation of Hog1 while the dissociation constant is kept constant.

### Modelling distributive or processive MAPK phosphorylation mechanisms in a basic three-tiered MAPK cascade

We utilized the paradigm of Huang and Ferrell, which models the basic three-tiered MAPK cascade. We collected or generated data measuring activation of MAP kinases from both yeast and mammalian cells that display different degrees of ultrasensitivity quantified by their Hill-coefficient. We found that Hog1 activation in *S. cerevisiae* by NaCl stimulation occurs with a hill-coefficient of 2.7, while its mammalian counterpart p38 in HeLa cells is stimulated by anisomycin with a hill-coefficient of 3 (Aoki et al., 2011). In comparison, the highly ultrasensitive activation of p38 via sorbitol in *Xenopus* oocytes occurs with a hill-coefficient of 14.4 (Ben Messaoud et al., 2015), and the graded response of Erk2 in HeLa cells stimulated by EGF with a hill-coefficient of 1.2 (Aoki et al., 2011).

The parameters of the basic model were optimized according to our described protocol and the best overall fit determined according to log-likelihood criteria. This was done for a distributive and a processive topology of MAPK activation. For Hog1 and both p38 data sets it was not possible to distinguish between a purely distributive or purely processive mechanism. Both models were able to fit the data equally well (difference in AIC was smaller than ten) and the utilized parameters still range within biologically feasible boundaries (Supp Figure S1C) (Bar-Even et al., 2011; Ben Messaoud et al., 2015). In contrast, the graded activation of Erk2 showed a clear preference for processive phosphorylation with significantly better fitting results, consistent with experimental evidence (Aoki et al., 2011). Experimental measurements also corroborate the possibility that mild ultrasensitive behavior can be achieved in nature using purely processive mechanisms. Thus, a processive mechanism with the correct parameters is also able to recapitulate even highly ultrasensitive behavior and thus cannot be readily discarded.

## Supplemental Information

**SI** Supp Figures S1-S5, Supp tables S1-S4

**Table S5** Model description: observables

**Table S6** Model description: volume sub-model

**Table S7** Model description: Sho1 sub-branch

**Table S8** Model description: Sln1 sub-branch

**Table S9** Model description: Hog1 activation

**Supp model files**

