## Supplementary material for "Positive feedback induces switch between distributive and processive phosphorylation of Hog1": Supp table S5

| Observables | Definition | Additional comment (respective position in Supp Figure 3) |
| --- | --- | --- |
| volume | (V_os + 1874)/(init_V_os + 1874) | Normalized change in volume (Cond1: volume; Cond5: volume; Cond6: volume; Cond7: volume; Cond8: volume) |
| hog1_ratio | (Hog1PPn + Hog1n + Hog1P174n + Hog1P176n)/(Hog1PPc + Hog1c + Hog1P174c + Hog1P176c + Hog1PPc_Pbs2PP + Hog1P174c_Pbs2PP + Hog1P176c_Pbs2PP + Hog1PPc_Pbs2PP_phosphorylated + Hog1P176c_Pbs2PP_phosphorylated + Hog1P174c_Pbs2PP_phosphorylated + Hog1c_Pbs2PP_phosphorylated + Hog1c_Pbs2 + Hog1P176c_Pbs2 + Hog1P174c_Pbs2 + Hog1PPc_Pbs2 + Hog1c_Pbs2_phosphorylated + Hog1P176c_Pbs2_phosphorylated + Hog1P174c_Pbs2_phosphorylated + Hog1PPc_Pbs2_phosphorylated) | Ratio between nuclear and cytosolic Hog1 (Cond1: Hog1 ratio; Cond5: Hog1 ratio; Cond6: Hog1 ratio; Cond7: Hog1 ratio; Cond8: Hog1 ratio; Cond13: Hog1 ratio) |
| output_hog1_ratio_mutant | See hog1_ratio | Ratio between nuclear and cytosolic Hog1 for data set with different error parameter (Cond2: Hog1 ratio mutant; Cond3: Hog1 ratio mutant; Cond4: Hog1 ratio mutant; Cond9: Hog1 ratio mutant; Cond10: Hog1 ratio mutant; Cond11: Hog1 ratio mutant; Cond12: Hog1 ratio mutant |
| output_Hog1PP | (molecules_Hog1PPc + molecules_Hog1PPn) /(starting_molecules_Hog1PPc + starting_molecules_Hog1PPn) | Relative ratio of double phosphorylated Hog1 (Cond1: Hog1PP 60s) |
| output_Hog1PP_vaga | See output_Hog1PP | Relative ratio of double phosphorylated Hog1 for data set with different error parameter (Cond1: Hog1PP) |
| Hog1_total_phosphorylation | (molecules_Hog1P174 + molecules_Hog1P176 + molecules_Hog1PPc + molecules_Hog1PPn)/(molecules_Hog1P174 + molecules_Hog1P176 + molecules_Hog1PPc + molecules_Hog1PPn +molecules_Hog1c + molecules_Hog1n)*100 | Percentage of mono or double phosphorylated Hog1 (Cond1: Hog1 total phosphorylation; Cond5: Hog1 total phosphorylation; Cond6: Hog1 total phosphorylation |
| Hog1_total_phosphorylation_inhibition | See Hog1_total_phosphorylation | Percentage of mono or double phosphorylated Hog1 for data sets with different error parameter (Cond14: Hog1 total phosphorylation; Cond15: Hog1 total parameter) |
| output_tyrosine_phosphorylation | ( molecules_Hog1P176 + molecules_Hog1PPc + molecules_Hog1PPn)/(molecules_Hog1P174 + molecules_Hog1P176 + molecules_Hog1PPc + molecules_Hog1PPn +molecules_Hog1c + molecules_Hog1n)*100 | Percentage with double phosphorylated or mono phosphorylated at tyrosine 176 Hog1 (Cond2: Hog1 tyrosine phosphorylation; Cond3: Hog1 tyrosine phosphorylation; Cond4: Hog1 tyrosine phosphorylation) |
| output_Pbs2P | (molecules_Pbs2P) / starting_molecules_Pbs2P | Relative rate of monophosphorylated Pbs2 (Cond1: Pbs2P 60s) |
| output_Hog1P174 | (molecules_Hog1P174) / starting_molecules_Hog1P174 | Relative ratio of Hog1 monophosphorylated at Threonine 174 (Cond1: Hog1P174 60s) |
| output_Hog1P174_vaga | See output_Hog1P174 | Relative ratio of Hog1 monophosphorylated at Threonine 174 for data set with different error parameter (Cond1: Hog1P174) |
| output_Hog1P176 | (molecules_Hog1P176) / starting_molecules_Hog1P176 | Relative ratio of Hog1 monophosphorylated at Tyrosine174 (Cond1: Hog1P176 60s) |
| output_Hog1P176_vaga | (molecules_Hog1P176) / starting_molecules_Hog1P176 | Relative ratio of Hog1 monophosphorylated at Tyrosine 174 for data set with different error parameter (Cond1: Hog1P176) |
| output_Ssk2_inactivation | (molecules_Ssk2_second_inactive )/(starting_molecules_Ssk2_second_inactive) | Relative ratio of double phosphorylated Ssk2 (Cond1: Ssk2PP feedback 60s) |
| output_Ssk2_inactivation_vaga | See output_Ssk2_inactivation | Relative ratio of double phosphorylated Ssk2 for data set with different error parameter (Cond1: Ssk2PP feedback) |
| output_Ssk2_inactivation_mono | (molecules_Ssk2_inactive )/(starting_molecules_Ssk2_inactive) | Relative ratio of mono phosphorylated Ssk2 (Cond1: Ssk2P feedback) |
| output_double_phosphorylation_ratio | (molecules_Hog1PPc + molecules_Hog1PPn)/(total_hog1) | Percentage of double phosphorylated Hog1 (Cond16: Hog1PP percentage) |
| output_Gpd1_ratio | (molecules_Gpd1 )/(starting_molecules_Gpd1) | Relative ratio of Gpd1 expression (Cond1: Gpd1) |
| output_Pbs2_phosphorylation_ratio | (molecules_Pbs2_phosphorylated)/ (starting_molecules_Pbs2_phosphorylated) | Relative ratio of Pbs2 phosphorylated by feedback (Cond1: Pbs2P feedback) |

| Species/Parameter | Definition | Additional comments |
| --- | --- | --- |
| to_molecule_number_cytosol | V_os_cytosol * 6.02214 | Conversion factor for concentration to absolute number of molecules in the cytosol |
| to_molecule_number_nucleus | V_os_nucleus * 6.02214 | Conversion factor for concentration to absolute number of molecules in the nucleus |
| to_molecule_number_cytosol_start | Init_V_os_cytosol * 6.02214 | Conversion factor for starting concentration to absolute number of molecules in the cytosol |
| to_molecule_number_nucleus_start | Init_V_os_nucleus * 6.02214 | Conversion factor for starting concentration to absolute number of molecules in the nucleus |
| molecules_Hog1PPc | (Hog1PPc + Hog1PPc_Pbs2PP + Hog1PPc_Pbs2PP_phosphorylated + Hog1PPc_Pbs2 + Hog1PPc_Pbs2_phosphorylated) * to_molecule_number_cytosol | Total number of double phosphorylated Hog1 in the cytosol |
| molecules_Hog1PPn | Hog1PPn*to_molecule_number_nucleus | Total number of double phosphorylated Hog1 in the nucleus |
| starting_molecules_Hog1PPc | init_Hog1PPc*to_molecule_number_cytosol_start | Total number of double phosphorylated Hog1 in the cytosol at the start |
| starting_molecules_Hog1PPn | init_Hog1PPn*to_molecule_number_nucleus_start | Total number of double phosphorylated Hog1 in the nucleus at the start |
| molecules_Hog1P174 | (Hog1P174c + Hog1P174c_Pbs2PP + Hog1P174c_Pbs2PP_phosphorylated + Hog1P174c_Pbs2 + Hog1P174c_Pbs2_phosphorylated ) * to_molecule_number_cytosol + Hog1P174n * to_molecule_number_nucleus | Total number of Hog1 monophosphorylated at Threonine 174 |
| molecules_Hog1P176 | (Hog1P176c + Hog1P176c_Pbs2PP + Hog1P176c_Pbs2PP_phosphorylated + Hog1P176c_Pbs2 + Hog1P176c_Pbs2_phosphorylated) * to_molecule_number_cytosol + Hog1P176n * to_molecule_number_nucleus | Total number of Hog1 monophosphorylated at Tyrosine 176 |
| molecules_Hog1c | (Hog1c + Hog1c_Pbs2PP + Hog1c_Pbs2PP_phosphorylated + Hog1c_Pbs2 + Hog1c_Pbs2_phosphorylated) * to_molecule_number_cytosol | Total number of unphosphorylated Hog1 in the cytosol |
| molecules_Hog1n | Hog1n*to_molecule_number_nucleus | Total number of unphosphorylated Hog1 in the nucleus |
| molecules_Pbs2P | (Pbs2P + Sho1_active_Ste11_Pbs2P + Pbs2P_phosphorylated + Sho1_active_Ste11_Pbs2P_phosphorylated) * to_molecule_number_cytosol | Total number of monophosphorylated Pbs2 |
| starting_molecules_Pbs2P | (init_Pbs2P + init_Pbs2P_phosphorylated)* to_molecule_number_cytosol_start | Total number of monophosphorylated Pbs2 at the start |
| molecules_Ssk2_second_inactive | (Ssk2_phosphorylated_second + Ssk2P_phosphorylated_second) * to_molecule_number_cytosol | Total number of double phosphorylated Ssk2 by feedback |
| molecules_Ssk2_inactive | (Ssk2_phosphorylated + Ssk2P_phosphorylated) * to_molecule_number_cytosol | Total number of mono phosphorylated Ssk2 by feedback |
| total_hog1 | (Hog1c + Hog1P174c + Hog1P176c + Hog1PPc + Hog1c_Pbs2PP + Hog1P174c_Pbs2PP + Hog1P176c_Pbs2PP + Hog1PPc_Pbs2PP + Hog1c_Pbs2_phosphorylated + Hog1P176c_Pbs2_phosphorylated + Hog1P174c_Pbs2_phosphorylated + Hog1PPc_Pbs2_phosphorylated + Hog1c_Pbs2PP_phosphorylated + Hog1P176c_Pbs2PP_phosphorylated + Hog1P174c_Pbs2PP_phosphorylated + Hog1PPc_Pbs2PP_phosphorylated + Hog1c_Pbs2 + Hog1P176c_Pbs2 + Hog1P174c_Pbs2 + Hog1PPc_Pbs2 )*to_molecule_number_cytosol + ( Hog1n + Hog1PPn + Hog1P174n + Hog1P176n)*to_molecule_number_nucleus | Total number of Hog1 of any phosphorylation state in the cell |
| molecules_Gpd1 | Gpd1 * to_molecule_number_cytosol | Total number of Gpd1 |
| molecules_Pbs2_phosphorylated | (Sho1_Pbs2_phosphorylated + Sho1_active_Pbs2_phosphorylated + Sho1_active_Ste11_Pbs2_phosphorylated_inactive + Sho1_active_Ste11_Pbs2_phosphorylated + Sho1_active_Ste11_Pbs2P_phosphorylated + Sho1_active_Ste11_Pbs2PP_phosphorylated + Sho1_active_Ste11_Pbs2_inactive_phosphorylated + Sho1_active_Ste11_Pbs2_phosphorylated + Sho1_active_Ste11_Pbs2P_phosphorylated + Pbs2_phosphorylated + Pbs2P_phosphorylated + Pbs2PP_phosphorylated + Hog1c_Pbs2_phosphorylated + Hog1P176c_Pbs2_phosphorylated + Hog1P174c_Pbs2_phosphorylated + Hog1PPc_Pbs2_phosphorylated + Hog1c_Pbs2PP_phosphorylated + Hog1P176c_Pbs2PP_phosphorylated + Hog1P174c_Pbs2PP_phosphorylated + Hog1PPc_Pbs2PP_phosphorylated) * to_molecule_number_cytosol | Total number of feedback phosphorylated Pbs2 |
| starting_molecules_Gpd1 | init_Gpd1 * to_molecule_number_cytosol_start | Total number of Gpd1 at the start |
| starting_molecules_Pbs2_phosphorylated | (init_Pbs2PP_phosphorylated + init_Pbs2P_phosphorylated + init_Pbs2_phosphorylated + init_Hog1c_Pbs2PP_phosphorylated + init_Hog1P174c_Pbs2PP_phosphorylated + init_Hog1P176c_Pbs2PP_phosphorylated + init_Hog1PPc_Pbs2PP_phosphorylated ) * to_molecule_number_cytosol_start | Total number of feedback phosphorylated Pbs2 at the start |
