## Supplementary material for "Positive feedback induces switch between distributive and processive phosphorylation of Hog1": Supp table S6

| **Glycerol production** | |
| --- | --- |
| Reaction | Reaction rate |
| -> glycerol | 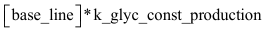 |
| glycerol -> | 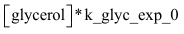 |
| glycerol -> | 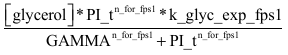 |
| Gpd1-> glycerol + Gpd1 | 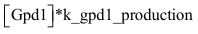 |
| -> glycerol | 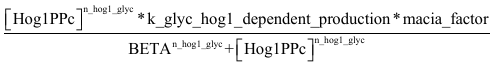 |
| Volume change and salt addition | |
| Reaction | Reaction rate |
| ->V_os_cytosol_ | 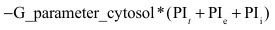 |
| ->V_os_nucleus_ | 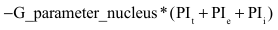 |
| ->V_os_ | 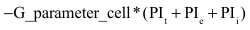 |
| ->NaCl | 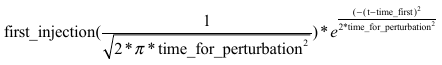 |

| **Algebraic pressure parameters** | |
| --- | --- |
| Parameter | Reaction rate |
| PI_t_ | 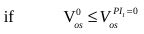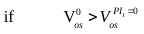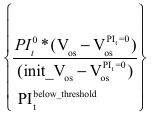 |
| PI_i_ | 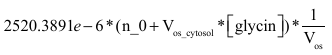 |
| PI_e_ | 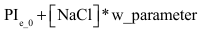 |

| Parameter | Parameter value^a^ | Unit | Additional information |
| --- | --- | --- | --- |
| k_glyc_const_production | 10.48 | s^-1^ | Constant basal production of glycerol |
| k_glyc_exp_0 | 3.98e-5 | s^-1^ | Rate of passive glycerol transport to the outside of the cell |
| n_for_fps1 | 9.92 | au |  |
| k_glyc_exp_fps1 | 0.09 | s^-1^ | Rate of glycerol transport to the outside via the Fps1 channel |
| GAMMA | 102.68 | 10^6^ J m^-3^ |  |
| k_gpd1_production | 100.00 | s^-1^ | Rate of glycerol production by Gpd1 |
| n_hog1_glyc | 7.14 | au |  |
| k_glyc_hog1_dependent_production | 43.83 | mM s^-1^ | Rate of glycerol production dependent on Hog1 activity in the cytosol |
| macia_factor | 1 | au | Factor that simulates inhibition of Hog1 activity when put to 0 |
| BETA | 0.03 | au |  |
| G_parameter_cytosol | 0.10 | 10^-24^m^6^J^-1^s^-1^ | Shrinkage factor for cytosol |
| G_parameter_nucleus | 0.02 | 10^-24^m^6^J^-1^s^-1^ | Shrinkage factor for nucleus |
| G_parameter_cell | 0.20 | 10^-24^m^6^J^-1^s^-1^ | Shrinkage factor for cell |
| first_injection | 400000 | μM | NaCl concentration added by a bolus injection |
| time_for_perturbation | 15 | s | Time duration of bolus injection |
| time_first | 2000 | s | Timepoint of NaCl injection |
| 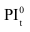 | 55.95 | 10^6^Jm^-3^ | Turgor pressure of unperturbed cell |
| 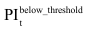 | 0.04 | 10^6^Jm^-3^ | Minimal turgor pressure |
| n_0 | 2.33e8 | 10^-15^μmol | Contribution by osmolarity other than glycerol |
| PI_e_0_ | 208.11 | 10^6^Jm^-3^ | External pressure in unperturbed state |
| w_parameter | 2.98e-4 | 10^6^Jm^-3^M^-1^ | Conversion factor of external salt concentration to external pressure |

| Species | Initial value (unit) | Additional information |
| --- | --- | --- |
| base_line | 1 (μM) | Proxy for the protein machinery needed for basal glycerol production |
| glycerol | 44218 (μM) | Intracellular glycerol |
| Hog1PPc | 0.0105 (mM) | Double phosphorylated Hog1 in the cytosol |
| V_os_cytosol_ | 2677 (10^-16^m^-3^) | Starting volume of the cytosol |
| V_os_nucleus_ | 267.7 (10^-16^m^-3^) | Starting volume of the nucleus |
| V_os_ | 3480 (10^-16^m^-3^) | Starting volume of the complete cell, includes volume that is considered incompressible and is not part of either cytosol or nucleus |
| NaCl | 1 (μM) | Extracellular NaCl concentration |

| **Gpd1 transcription** | |
| --- | --- |
| Reaction | Reaction rate |
| Gene_off -> Gene_on | 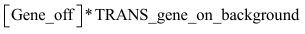 |
| Gene_on -> Gene_off | 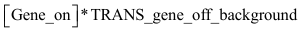 |
| Gene_off + Hog1PPn -> Gene_on_Hog1PPn | 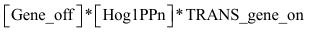 |
| Gene_on_Hog1PPn -> Gene_off + Hog1PPn | 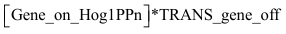 |
| Gene_on -> Gene_on_r | 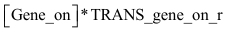 |
| Gene_on_Hog1PPn -> Gene_on_r_Hog1PPn | 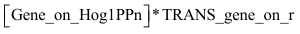 |
| Gene_on_r -> Gene_on | 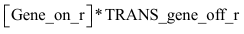 |
| Gene_on_r_Hog1PPn -> Gene_on_Hog1PPn | 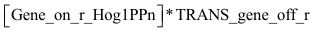 |
| Gene_on_r -> Gene_on_r + mRNA | 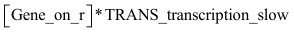 |
| Gene_on_r_Hog1PPn -> Gene_on_r_Hog1PPn + mRNA | 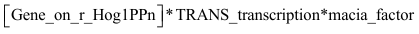 |
| mRNA -> | 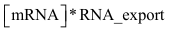 |
| -> mRNA_cytosol | 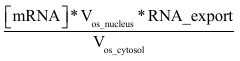 |
| mRNA_cytosol -> mRNA_cytosol + Gpd1_pre | 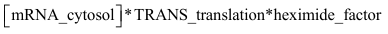 |
| mRNA_cytosol -> | 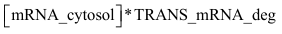 |
| Gpd1_pre -> Gpd1 | 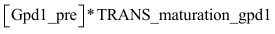 |
| Gpd1 -> | 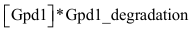 |

| Parameter | Parameter value^a^ | Unit | Additional information |
| --- | --- | --- | --- |
| TRANS_gene_on_background | 1.4887 | s^-1^ | Slow, basal rate of closed complex formation at promoter for transcription |
| TRANS_gene_off_background | 0.2462 | s^-1^ | Basal rate of disassociation of closed complex from promoter (gene shut off) |
| TRANS_gene_on | 83.4449 | mM^-1^s^-1^ | Rate of priming of promoter for transcription with double phosphorylated Hog1 |
| TRANS_gene_off | 0.1892 | s^-1^ | Rate of disassociation of closed complex from promoter (gene shut off) with double phosphorylated Hog1 bound |
| TRANS_Gene_on_r | 0.0011 | s^-1^ | Rate of formation of open complex |
| TRANS_gene_off_r | 0.0041 | s^-1^ | Rate of closure of open complex |
| TRANS_transcription_slow | 0.0554 | s^-1^ | Slow, basal transcription rate |
| TRANS_transcription | 15.6135 | s^-1^ | Transcription rate with double phosphorylated Hog1 bound |
| macia_factor | 1 | au | Factor that simulates inhibition of Hog1 activity when put to 0 preventing double phosphorylated Hog1 to bind to DNA |
| RNA_export | 55.6801 | s^-1^ | Rate of RNA export into the cytosol |
| V_os_cytosol_ | See above |  | See Volume sub-model |
| V_os_nucleus_ | See above |  | See Volume sub-model |
| TRANS_translation | 1.3533 | s^-1^ | Rate of translation |
| Heximide_factor | 1 | au | Factor that simulates inhibition of translational activity when put to 0 such as by the addition of Heximide |
| TRANS_maturation_gpd1 | 9.7949e-04 | s^-1^ | Rate of protein folding of Gpd1 |
| Gpd1_degradation | 3.7574e-04 | s^-1^ | Rate of degradation of the Gpd1 protein |

| Species | Initial value (unit) | Additional information |
| --- | --- | --- |
| Gene_off | 1.0000e-04 (mM) | Arbitrary concentration value that represents Gene in off state |
| Gene_on | 0 (mM) | Gene in on state |
| Hog1PPn | 0 (mM) | Concentration of double phosphorylated Hog1 in the nucleus |
| Gene_on_r | 0 (mM) | Gene with transcription complex in open position |
| Gene_on_Hog1PPn | 0 (mM) | Double phosphorylated Hog1 bound to Gene in closed position |
| Gene_on_r_Hog1PPn | 0 (mM) | Double phosphorylated Hog1 bound to Gene in open position |
| mRNA | 0.0025 (mM) | Concentration of mRNA in the nucleus |
| mRNA_cytosol | 0 (mM) | Concentration of mRNA in the cytosol |
| Gpd1_pre | 0 (mM) | Concentration of unfolded Gpd1 protein |
| Gpd1_a_ | 0.05 (mM) | Concentration of folded Gpd1 |

_a_ Total amount of Gpd1 in the cell: 807 molecules[1]

1.	Ghaemmaghami, S., et al., *Global analysis of protein expression in yeast.* Nature, 2003. **425**(6959): p. 737-41.
