## Supplementary material for "Positive feedback induces switch between distributive and processive phosphorylation of Hog1": Supp table S9

| **Hog1 shuttling** | |
| --- | --- |
| Reaction | Reaction rate |
| Hog1c -> | 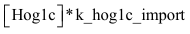 |
| Hog1n -> | 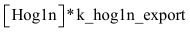 |
| -> Hog1n | 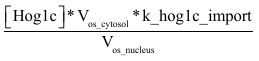 |
| -> Hog1c | 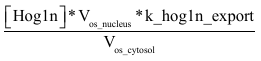 |
| Hog1PPc -> | 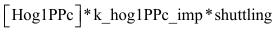 |
| Hog1PPn -> | 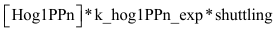 |
| -> Hog1PPn | 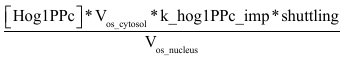 |
| -> Hog1PPc | 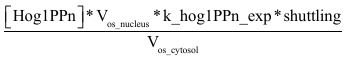 |
| Hog1P176c -> | 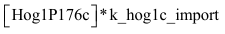 |
| Hog1P176n -> | 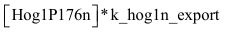 |
| -> Hog1P176n | 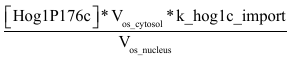 |
| -> Hog1P176c | 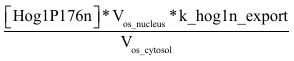 |
| Hog1P174c -> | 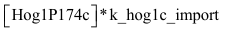 |
| Hog1P174n -> | 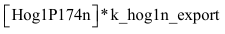 |
| -> Hog1P174n | 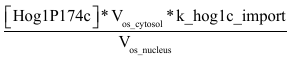 |
| -> Hog1P174c | 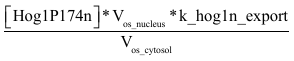 |

| Parameter | Parameter value^a^ | Unit | Additional information |
| --- | --- | --- | --- |
| k_hog1c_import | 0.1346 | s^-1^ | Passive, basal import of non-phosphorylated and mono-phosphorylated Hog1 |
| k_hog1n_export | 0.8902 | s^-1^ | Passive, basal export of non-phosphorylated and mono-phosphorylated Hog1 |
| V_os_cytosol_ | var |  | Variable parameter of the changing cytosol volume depending on pressure: see Supp Volume sub-model |
| V_os_nucleus_ | var |  | Variable parameter of the changing nuclear volume depending on pressure: see Supp Volume sub-model |
| K_hog1PPc_imp | 0.0231 | s^-1^ | Active import of double phosphorylated Hog1 |
| K_hog1PPn_exp | 1.3125e-6 | s^-1^ | Active export of double phosphorylated Hog1 is basically non-existent |
| shuttling | 1 | dimensionless | Auxilliary parameter to switch shuttling on or off |

| **Hog1 phosphorylation mixed mechanism** | |
| --- | --- |
| Hog1c + Pbs2PP -> Hog1c_Pbs2PP | 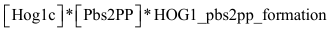 |
| Hog1c_Pbs2PP -> Hog1c + Pbs2PP | 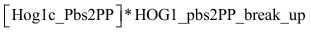 |
| Hog1c_Pbs2PP -> Hog1P174c_Pbs2PP | 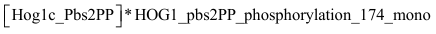 |
| Hog1c_Pbs2PP -> Hog1P176c_Pbs2PP | 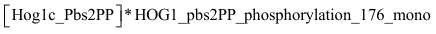 |
| Hog1P176c + Pbs2PP -> Hog1P176c_Pbs2PP | 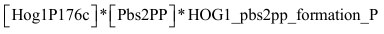 |
| Hog1P174c + Pbs2PP -> Hog1P174c_Pbs2PP | 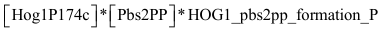 |
| Hog1P174c_Pbs2PP -> Hog1P174c + Pbs2PP | 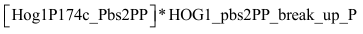 |
| Hog1P176c_Pbs2PP -> Hog1P176c + Pbs2PP | 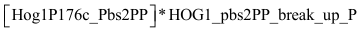 |
| Hog1P174c_Pbs2PP -> Hog1PPc_Pbs2PP | 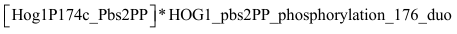 |
| Hog1P176c_Pbs2PP -> Hog1PPc_Pbs2PP | 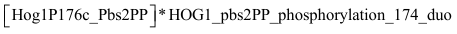 |
| Hog1PPc + Pbs2PP -> Hog1PPc_Pbs2PP | 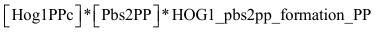 |
| Hog1PPc_Pbs2PP -> Hog1PPc + Pbs2PP | 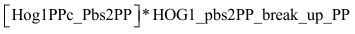 |
| Hog1c + Pbs2PP_phosphorylated -> Hog1c_Pbs2PP_phosphorylated | 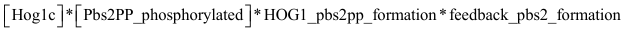 |
| Hog1c_Pbs2PP_phosphorylated -> Hog1c + Pbs2PP_phosphorylated | 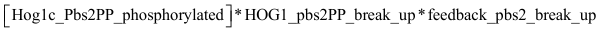 |
| Hog1c_Pbs2PP_phosphorylated -> Hog1P174c_Pbs2PP_phosphorylated | 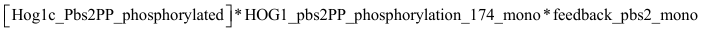 |
| Hog1c_Pbs2PP_phosphorylated -> Hog1P176c_Pbs2PP_phosphorylated | 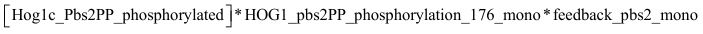 |
| Hog1P176c + Pbs2PP_phosphorylated -> Hog1P176c_Pbs2PP_phosphorylated | 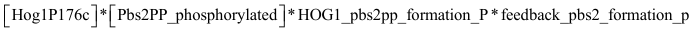 |
| Hog1P174c + Pbs2PP_phosphorylated -> Hog1P174c_Pbs2PP_phosphorylated | 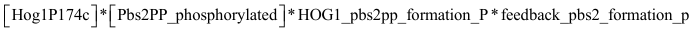 |
| Hog1P174c_Pbs2PP_phosphorylated -> Hog1P174c + Pbs2PP_phosphorylated | 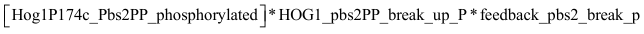 |
| Hog1P176c_Pbs2PP_phosphorylated -> Hog1P176c + Pbs2PP_phosphorylated | 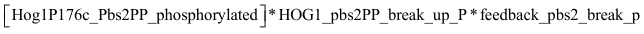 |
| Hog1P174c_Pbs2PP_phosphorylated -> Hog1PPc_Pbs2PP_phosphorylated | 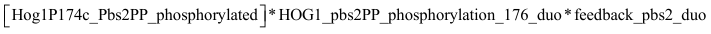 |
| Hog1P176c_Pbs2PP_phosphorylated -> Hog1PPc_Pbs2PP_phosphorylated | 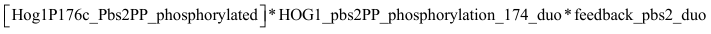 |
| Hog1PPc + Pbs2PP_phosphorylated -> Hog1PPc_Pbs2PP_phosphorylated | 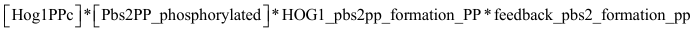 |
| Hog1PPc_Pbs2PP_phosphorylated -> Hog1PPc + Pbs2PP_phosphorylated | 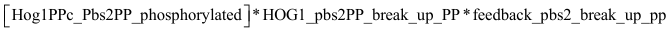 |
| **Formation of Hog1 – Pbs2 complexes that act as targets for the Ptc1 phosphatase** | |
| Hog1c + Pbs2 -> Hog1c_Pbs2 | 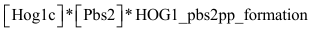 |
| Hog1c_Pbs2 -> Hog1c + Pbs2 | 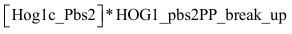 |
| Hog1P176c + Pbs2 -> Hog1P176c_Pbs2 | 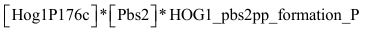 |
| Hog1P174c + Pbs2 -> Hog1P174c_Pbs2 | 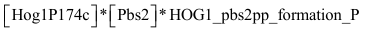 |
| Hog1P174c_Pbs2 -> Hog1P174c + Pbs2 | 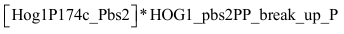 |
| Hog1P176c_Pbs2 -> Hog1P176c + Pbs2 | 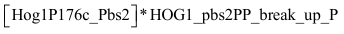 |
| Hog1PPc + Pbs2 -> Hog1PPc_Pbs2 | 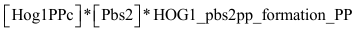 |
| Hog1PPc_Pbs2 -> Hog1PPc + Pbs2 | 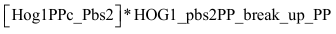 |
| **Formation of Hog1 – Pbs2 with feedback complexes that act as targets for the Ptc1 phosphatase** | |
| Hog1c + Pbs2_phosphorylated -> Hog1c_Pbs2_phosphorylated | 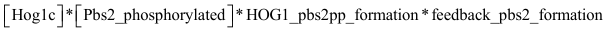 |
| Hog1c_Pbs2_phosphorylated -> Hog1c + Pbs2_phosphorylated | 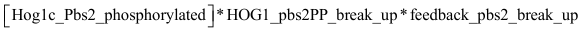 |
| Hog1P176c + Pbs2_phosphorylated -> Hog1P176c_Pbs2_phosphorylated | 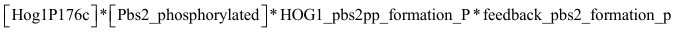 |
| Hog1P174c + Pbs2_phosphorylated -> Hog1P174c_Pbs2_phosphorylated | 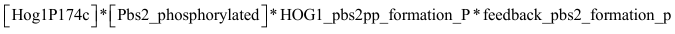 |
| Hog1P174c_Pbs2_phosphorylated -> Hog1P174c + Pbs2_phosphorylated | 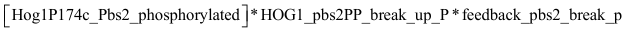 |
| Hog1P176c_Pbs2_phosphorylated -> Hog1P176c + Pbs2_phosphorylated | 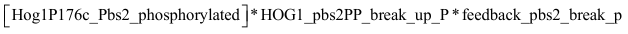 |
| Hog1PPc + Pbs2_phosphorylated -> Hog1PPc_Pbs2_phosphorylated | 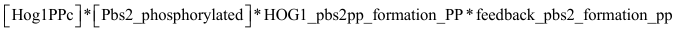 |
| Hog1PPc_Pbs2_phosphorylated -> Hog1PPc + Pbs2_phosphorylated | 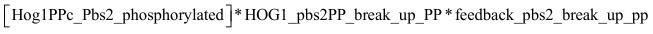 |

| Parameter | Parameter value^a^ | Unit | Additional information |
| --- | --- | --- | --- |
| HOG1_pbs2pp_formation | 153.85 | mM^-1^s^-1^ | Rate of association between Hog1 and Pbs2PP |
| HOG1_pbs2pp_break_up | 921.51 | s^-1^ | Rate of disassociation between Hog1 and Pbs2PP |
| HOG1_pbs2pp_formation_P | 60.39 | mM^-1^s^-1^ | Rate of association between monophosphorylated Hog1 and Pbs2PP |
| HOG1_pbs2pp_break_up_P | 828.51 | s^-1^ | Rate of disassociation between monophosphorylated Hog1 and Pbs2PP |
| HOG1_pbs2pp_formation_PP | 180.55 | mM^-1^s^-1^ | Rate of association between double phosphorylated Hog1 and Pbs2PP |
| HOG1_pbs2pp_break_up_PP | 56.34 | s^-1^ | Rate of disassociation between double phosphorylated Hog1 and Pbs2PP |
| HOG1_pbs2pp_phosphorylation_174_duo | 4.23 | s^-1^ | Phosphorylation of Thr174 on Hog1P176 |
| HOG1_pbs2pp_phosphorylation_174_mono | 71.17 | s^-1^ | Phosphorylation of Thr174 on non-phosphorylated Hog1 |
| HOG1_pbs2pp_phosphorylation_176_duo | 1.94 | s^-1^ | Phosphorylation of Tyr176 on Hog1P174 |
| HOG1_pbs2pp_phosphorylation_176_mono | 290.54 | s^-1^ | Phosphorylation of Tyr176 on non-phosphorylated Hog1 |
| feedback_pbs2_formation | 0.89 | dimensionless | Change in the association rate between Hog1 and Pbs2 |
| feedback_pbs2_break_up | 6.18 | dimensionless | Change in the disassociation rate between Hog1 and Pbs2 |
| feedback_pbs2_formation_p | 0.18 | dimensionless | Change in the association rate between Hog1P and Pbs2 |
| feedback_pbs2_break_p | 0.16 | dimensionless | Change in the disassociation rate between Hog1P and Pbs2 |
| feedback_pbs2_formation_pp | 1.67 | dimensionless | Change in the association rate between Hog1PP and Pbs2 |
| feedback_pbs2_break_up_pp | 5.52 | dimensionless | Change in the disassociation rate between Hog1 and Pbs2 |
| feedback_pbs2_mono | 2.40 | dimensionless | Change in the phosphorylation rate of the first phosphorylation of Hog1 |
| feedback_pbs2_duo | 66.70 | dimensionless | Change in the phosphorylation rate of the second phosphorylation of Hog1 |

| **Hog1 dephosphorylation** | |
| --- | --- |
| Hog1PPc + additional_phosphatase_cyto -> Hog1P174c + additional_phosphatase_cyto | 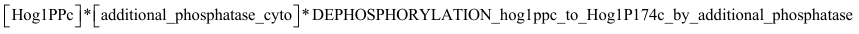 |
| Hog1PPn + additional_phosphatase_nucleus -> Hog1P174n + additional_phosphatase_nucleus | 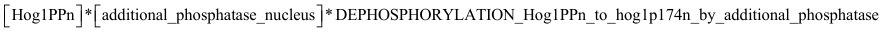 |
| Hog1P176c + additional_phosphatase_cyto -> Hog1c + additional_phosphatase_cyto | 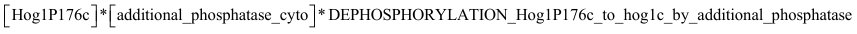 |
| Hog1P176n + additional_phosphatase_nucleus -> Hog1n + additional_phosphatase_nucleus | 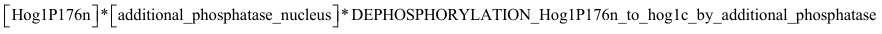 |
| Hog1PPc + Ptp3 -> Hog1P174c + Ptp3 | 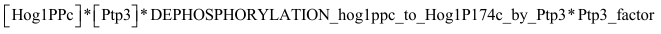 |
| Hog1P176c + Ptp3 -> Hog1c + Ptp3 | 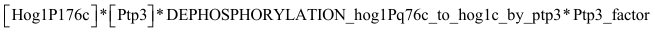 |
| Hog1PPc_Pbs2PP + Ptc1 -> Hog1P176c_Pbs2PP + Ptc1 | 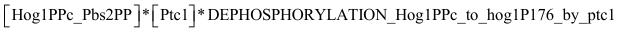 |
| Hog1PPc_Pbs2 + Ptc1 -> Hog1P176c_Pbs2 + Ptc1 | 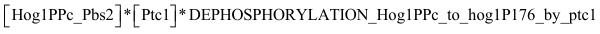 |
| Hog1PPc_Pbs2PP_phosphorylated + Ptc1 -> Hog1P176c_Pbs2PP_phosphorylated + Ptc1 | 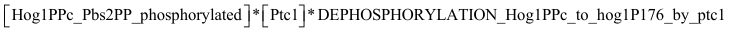 |
| Hog1PPc_Pbs2_phosphorylated + Ptc1 -> Hog1P176c_Pbs2_phosphorylated + Ptc1 | 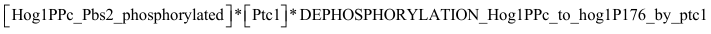 |
| Hog1P174c_Pbs2PP + Ptc1 -> Hog1c_Pbs2PP + Ptc1 | 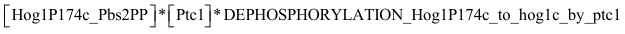 |
| Hog1P174c_Pbs2 + Ptc1 -> Hog1c_Pbs2 + Ptc1 | 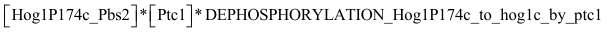 |
| Hog1P174c_Pbs2PP_phosphorylated + Ptc1 -> Hog1c_Pbs2PP_phosphorylated + Ptc1 | 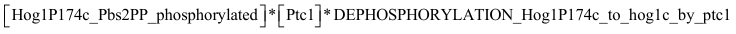 |
| Hog1P174c_Pbs2_phosphorylated + Ptc1 -> Hog1c_Pbs2_phosphorylated + Ptc1 | 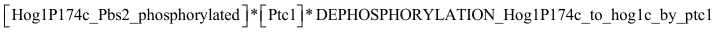 |
| Hog1PPn + Ptp2 -> Hog1P174n + Ptp2 | 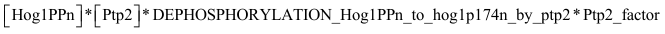 |
| Hog1PPn + Ptc -> Hog1P176n + Ptc | 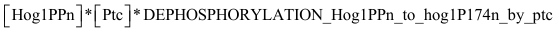 |
| Hog1P174n + Ptc -> Hog1n + Ptc | 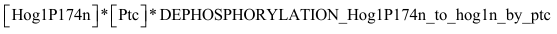 |
| Hog1P176n + Ptp2 -> Hog1n + Ptp2 | 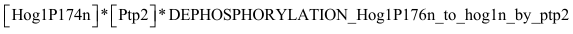 |

| Parameter | Parameter value^a^ | Unit | Additional information |
| --- | --- | --- | --- |
| DEPHOSPHORYLATION_Hog1PPn_to_hog1p174n_by_additional_phosphatase | 0.0019 | mM^-1^s^-1^ | Dephosphorylation by unspecific phosphatases |
| DEPHOSPHORYLATION_hog1ppc_to_Hog1P174c_by_additional_phosphatase | 2e-19 | mM^-1^s^-1^ | Dephosphorylation by unspecific phosphatases |
| DEPHOSPHORYLATION_Hog1P176c_to_hog1c_by_additional_phosphatase | 3.68e-8 | mM^-1^s^-1^ | Dephosphorylation by unspecific phosphatases |
| DEPHOSPHORYLATION_Hog1P176n_to_hog1n_by_additional_phosphatase | 0.0133 | mM^-1^s^-1^ | Dephosphorylation by unspecific phosphatases |
| DEPHOSPHORYLATION_hog1ppc_to_Hog1P174c_by_Ptp3 | 7.3621e-17 | mM^-1^s^-1^ | Dephosphorylation of Tyrosine in the cytosol |
| DEPHOSPHORYLATION_Hog1P176c_to_hog1c_by_ptp3 | 0.0019 | mM^-1^s^-1^ | Dephosphorylation of Tyrosine in the cytosol |
| DEPHOSPHORYLATION_Hog1PPn_to_hog1p174n_by_ptp2 | 2.18e-5 | mM^-1^s^-1^ | Dephosphorylation of Tyrosine in the nucleus |
| DEPHOSPHORYLATION_Hog1P176n_to_hog1n_by_ptp2 | 2.0735 | mM^-1^s^-1^ | Dephosphorylation of Tyrosine in the cytosol |
| DEPHOSPHORYLATION_Hog1P174c_to_hog1c_by_ptc1 | 2.3318 | s^-1^ | Dephosphorylation by Ptc1 when bound to Pbs2 |
| DEPHOSPHORYLATION_Hog1PPc_to_hog1P176_by_ptc1 | 5.46e-4 | s^-1^ | Dephosphorylation by Ptc1 when bound to Pbs2 |
| DEPHOSPHORYLATION_Hog1P174n_to_hog1n_by_ptc | 0.5077 | mM^-1^s^-1^ | Dephosphorylation by Ptc2/3 in Nucleus |
| DEPHOSPHORYLATION_Hog1PPn_to_hog1P176n_by_ptc | 0.0563 | mM^-1^s^-1^ | Dephosphorylation by Ptc2/3 in Nucleus |

| Species | Initial value (unit) | Additional information |
| --- | --- | --- |
| Hog1c_a_ | 0.3109 (mM) | Cytosolic Hog1 |
| Hog1n | 0.5278 (mM) | Nuclear Hog1 |
| Hog1PPc | 0.0105 (mM) | Double phosphorylated, cytosolic Hog1 |
| Hog1PPn | 0 (mM) | Double phosphorylated, nuclear Hog1 |
| Hog1P176c | 0.0303 (mM) | Cytosolic Hog1 phosphorylated at Tyr-176 |
| Hog1P176n | 0 (mM) | Nuclear Hog1 phosphorylated at Tyr-176 |
| Hog1P174c | 0.0026 (mM) | Cytosolic Hog1 phosphorylated at Thr-174 |
| Hog1P174n | 0 (mM) | Nuclear Hog1 phosphorylated at Thr-174 |
| Pbs2PP | 0 (mM) | activated, double phosphorylated Pbs2 |
| Hog1c_Pbs2PP | 0 (mM) | Hog1 bound to activated, double phosphorylated Pbs2 |
| Hog1P174c_Pbs2PP | 0 (mM) | Hog1 phosphorylated at Thr-174 bound to activated, double phosphorylated Pbs2 |
| Hog1P176c_Pbs2PP | 0 (mM) | Hog1 phosphorylated at Tyr-176 bound to activated, double phosphorylated Pbs2 |
| Hog1PPc_Pbs2PP | 0 (mM) | Double phosphorylated Hog1 bound to activated, double phosphorylated Pbs2 |
| Pbs2PP_phosphorylated | 0 (mM) | activated, double phosphorylated Pbs2 with additional feedback phosphorylation |
| Hog1c_Pbs2PP_phosphorylated | 0 (mM) | Hog1 bound to activated, double phosphorylated Pbs2 with additional feedback phosphorylation |
| Hog1P174c_Pbs2PP_phosphorylated | 0 (mM) | Hog1 phosphorylated at Thr-174 bound to activated, double phosphorylated Pbs2 with additional feedback phosphorylation |
| Hog1P176c_Pbs2PP_phosphorylated | 0 (mM) | Hog1 phosphorylated at Tyr-176 bound to activated, double phosphorylated Pbs2 with additional feedback phosphorylation |
| Hog1PPc_Pbs2PP_phosphorylated | 0 (mM) | Double phosphorylated Hog1 bound to activated, double phosphorylated Pbs2 with additional feedback phosphorylation |
| Pbs2_b_ | 0.1138 (mM) | MAPKK of Hog1 |
| Hog1P176c_Pbs2 | 0 (mM) | Hog1 phosphorylated at Tyr-176 bound to Pbs2 |
| Hog1P174c_Pbs2 | 0 (mM) | Hog1 phosphorylated at Thr-174 bound to Pbs2 |
| Hog1PPc_Pbs2 | 0 (mM) | Double phosphorylated Hog1 bound to Pbs2 |
| Pbs2_phosphorylated | 0.01 (mM) | Pbs2 with additional feedback phosphorylation |
| Pbs2PP_phosphorylated | 1.26e-6 (mM) | activated, double phosphorylated Pbs2 with additional feedback phosphorylation |
| Hog1P176c_Pbs2_phosphorylated | 0 (mM) | Hog1 phosphorylated at Tyr-176 bound to Pbs2 with additional feedback phosphorylation |
| Hog1P174c_Pbs2_phosphorylated | 0 (mM) | Hog1 phosphorylated at Thr-174 bound to Pbs2 with additional feedback phosphorylation |
| Hog1PPc_Pbs2_phosphorylated | 0 (mM) | Double phosphorylated Hog1 bound to Pbs2 with additional feedback phosphorylation |
| additional_phosphatase_cyto | 1 (mM) | Arbitrary approximation of all unspecific phosphatases in the cytosol |
| additional_phosphatase_nucleus | 1 (mM) | Arbitrary approximation of all unspecific phosphatases in the nucleus |
| Ptp3_c_ | 0.48 (mM) | Cytosolic Ptp3 |
| Ptc1_d_ | 0.0943 (mM) | Cytosolic Ptc1 |
| Ptp2_e_ | 0.089 (mM) | Nuclear Ptp2 |
| Ptc_f_ | 1.3576 (mM) | Nuclear proportion of Ptc2/3 when molecules are evenly distributed throughout cell |

_a_ Total amount of Hog1 in the cell: 6780 molecules[1]

_b_ Total amount of Pbs2 in the cell: 2160 molecules[1]

_c_ Total amount of Ptp3 in the cell: 768 molecules[1]

_d_ Total amount of Ptc1 in the cell: 1520 molecules[1]

_e_ Total amount of Ptp2 in the cell: 149 molecules[1]

_f_ Total amount of Ptc2/3 in the nucleus: 7% of 12600 molecules of Ptc2[1] and 19700 molecules of Ptc3[1]

1.	Ghaemmaghami, S., et al., *Global analysis of protein expression in yeast.* Nature, 2003. **425**(6959): p. 737-41.

**Reactions used for the distributive mechanism of Hog1 phosphorylation**

| **Hog1 phosphorylation distributive mechanism** | |
| --- | --- |
| Hog1c + Pbs2PP -> Hog1c_Pbs2PP | 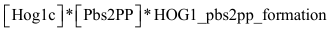 |
| Hog1c_Pbs2PP -> Hog1c + Pbs2PP | 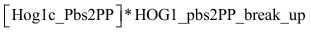 |
| Hog1c_Pbs2PP -> Hog1P174c + Pbs2PP | 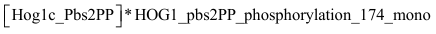 |
| Hog1c_Pbs2PP -> Hog1P176c + Pbs2PP | 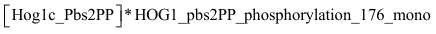 |
| Hog1P176c + Pbs2PP -> Hog1P176c_Pbs2PP | 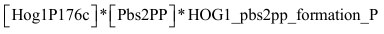 |
| Hog1P174c + Pbs2PP -> Hog1P174c_Pbs2PP | 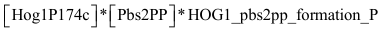 |
| Hog1P174c_Pbs2PP -> Hog1P174c + Pbs2PP | 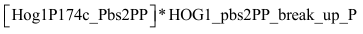 |
| Hog1P176c_Pbs2PP -> Hog1P176c + Pbs2PP | 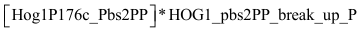 |
| Hog1P174c_Pbs2PP -> Hog1PPc_Pbs2PP | 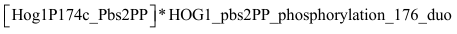 |
| Hog1P176c_Pbs2PP -> Hog1PPc_Pbs2PP | 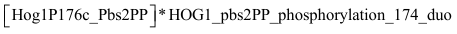 |
| Hog1PPc + Pbs2PP -> Hog1PPc_Pbs2PP | 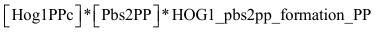 |
| Hog1PPc_Pbs2PP -> Hog1PPc + Pbs2PP | 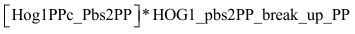 |
| Hog1c + Pbs2PP_phosphorylated -> Hog1c_Pbs2PP_phosphorylated | 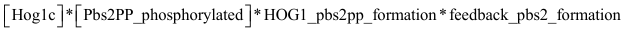 |
| Hog1c_Pbs2PP_phosphorylated -> Hog1c + Pbs2PP_phosphorylated | 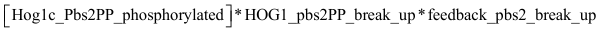 |
| Hog1c_Pbs2PP_phosphorylated -> Hog1P174c + Pbs2PP_phosphorylated | 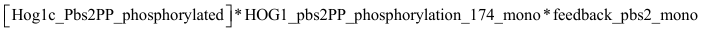 |
| Hog1c_Pbs2PP_phosphorylated -> Hog1P176c + Pbs2PP_phosphorylated | 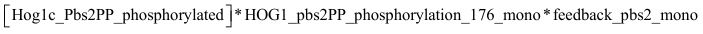 |
| Hog1P176c + Pbs2PP_phosphorylated -> Hog1P176c_Pbs2PP_phosphorylated | 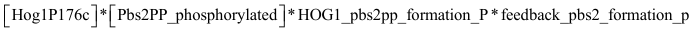 |
| Hog1P174c + Pbs2PP_phosphorylated -> Hog1P174c_Pbs2PP_phosphorylated | 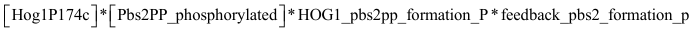 |
| Hog1P174c_Pbs2PP_phosphorylated -> Hog1P174c + Pbs2PP_phosphorylated | 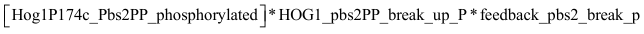 |
| Hog1P176c_Pbs2PP_phosphorylated -> Hog1P176c + Pbs2PP_phosphorylated | 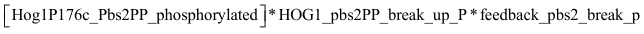 |
| Hog1P174c_Pbs2PP_phosphorylated -> Hog1PPc_Pbs2PP_phosphorylated | 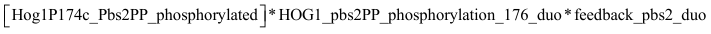 |
| Hog1P176c_Pbs2PP_phosphorylated -> Hog1PPc_Pbs2PP_phosphorylated | 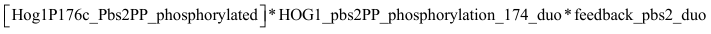 |
| Hog1PPc + Pbs2PP_phosphorylated -> Hog1PPc_Pbs2PP_phosphorylated | 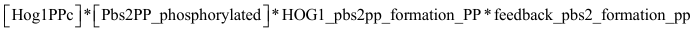 |
| Hog1PPc_Pbs2PP_phosphorylated -> Hog1PPc + Pbs2PP_phosphorylated | 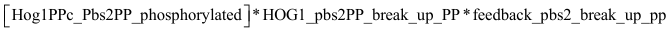 |

**Reactions used for the processive mechanism of Hog1 phosphorylation (reactions highlighted in red have been removed)**

| **Hog1 phosphorylation processive mechanism** | |
| --- | --- |
| Hog1c + Pbs2PP -> Hog1c_Pbs2PP | 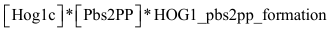 |
| Hog1c_Pbs2PP -> Hog1c + Pbs2PP |  |
| Hog1c_Pbs2PP -> Hog1P174c_Pbs2PP |  |
| Hog1c_Pbs2PP -> Hog1P176c_Pbs2PP |  |
| Hog1P176c + Pbs2PP -> Hog1P176c_Pbs2PP |  |
| Hog1P174c + Pbs2PP -> Hog1P174c_Pbs2PP |  |
| Hog1P174c_Pbs2PP -> Hog1P174c + Pbs2PP |  |
| Hog1P176c_Pbs2PP -> Hog1P176c + Pbs2PP |  |
| Hog1P174c_Pbs2PP -> Hog1PPc_Pbs2PP |  |
| Hog1P176c_Pbs2PP -> Hog1PPc_Pbs2PP |  |
| Hog1PPc + Pbs2PP -> Hog1PPc_Pbs2PP |  |
| Hog1PPc_Pbs2PP -> Hog1PPc + Pbs2PP |  |
| Hog1c + Pbs2PP_phosphorylated -> Hog1c_Pbs2PP_phosphorylated |  |
| Hog1c_Pbs2PP_phosphorylated -> Hog1c + Pbs2PP_phosphorylated |  |
| Hog1c_Pbs2PP_phosphorylated -> Hog1P174c_Pbs2PP_phosphorylated |  |
| Hog1c_Pbs2PP_phosphorylated -> Hog1P176c_Pbs2PP_phosphorylated |  |
| Hog1P176c + Pbs2PP_phosphorylated -> Hog1P176c_Pbs2PP_phosphorylated |  |
| Hog1P174c + Pbs2PP_phosphorylated -> Hog1P174c_Pbs2PP_phosphorylated |  |
| Hog1P174c_Pbs2PP_phosphorylated -> Hog1P174c + Pbs2PP_phosphorylated |  |
| Hog1P176c_Pbs2PP_phosphorylated -> Hog1P176c + Pbs2PP_phosphorylated |  |
| Hog1P174c_Pbs2PP_phosphorylated -> Hog1PPc_Pbs2PP_phosphorylated |  |
| Hog1P176c_Pbs2PP_phosphorylated -> Hog1PPc_Pbs2PP_phosphorylated |  |
| Hog1PPc + Pbs2PP_phosphorylated -> Hog1PPc_Pbs2PP_phosphorylated |  |
| Hog1PPc_Pbs2PP_phosphorylated -> Hog1PPc + Pbs2PP_phosphorylated |  |
