## Supplementary material for "Positive feedback induces switch between distributive and processive phosphorylation of Hog1": Supp table S8

| **Sln1 sub-branch** | |
| --- | --- |
| Reaction | Reaction rate |
| Sln1 -> Sln1P | 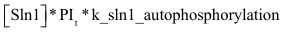 |
| Sln1P + Ypd1-> Sln1 + Ypd1P | 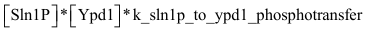 |
| Sln1 + Ypd1P -> Sln1P + Ypd1 | 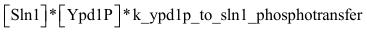 |
| Ypd1P + Ssk1 -> Ypd1 + Ssk1P | 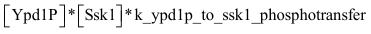 |
| Ssk1P -> Ssk1 | 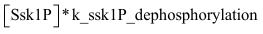 |
| Ssk2 + Ssk1 -> Ssk2P + Ssk1 | 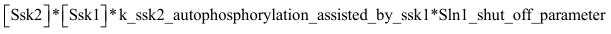 |
| Ssk2P -> Ssk2 | 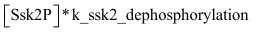 |
| Ssk2P + Pbs2 -> Pbs2P + Ssk2P | 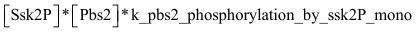 |
| Pbs2P + Ssk2P -> Pbs2PP + Ssk2P | 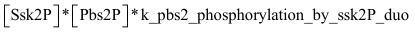 |
| Pbs2PP -> Pbs2P | 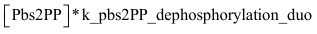 |
| Pbs2P -> Pbs2 | 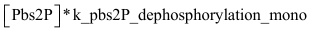 |

| Parameter | Parameter value | Unit | Additional information |
| --- | --- | --- | --- |
| PI_t_ | variable |  | Variable turgor pressure. See Volume sub-model |
| k_sln1_autophosphorylation | 0.01 | s^-1^ | Basal autophosphorylation of Sln1 |
| k_sln1p_to_ypd1_phosphotransfer | 51.92 | mM^-1^s^-1^ | Transfer of phosphate from Sln1 to Ypd1 |
| k_ypd1P_to_sln1_phosphotransfer | 418.02 | mM^-1^s^-1^ | Transfer of phosphate from Ypd1 to Sln1 |
| k_ypd1p_to_ssk1_phosphotransfer | 999.77 | mM^-1^s^-1^ | Transfer of phosphate from Ypd1 to Ssk1 |
| k_ssk1P_dephosphorylation | 0.23 | s^-1^ | Dephosphorylation of Ssk1 by unspecified phosphatase |
| k_ssk2_autophosphorylation_assisted_by_ssk1 | 28.61 | mM^-1^s^-1^ | Autophosphorylation of Ssk2 induced by association to Ssk1 |
| k_ssk2_dephosphorylation | 1.38 | s^-1^ | Dephosphorylation of Ssk2 by unspecified phosphatase |
| k_pbs2_phosphorylation_by_ssk2P_mono | 448.75 | mM^-1^s^-1^ | Phosphorylation of Pbs2 by phosphorylated Ssk2 |
| k_pbs2p_phosphorylation_by_ssk2P_duo | 434.41 | mM^-1^s^-1^ | Phosphorylation of Pbs2P by phosphorylated Ssk2 |
| k_pbs2PP_dephosphorylation_duo | 0.83 | s^-1^ | Dephosphorylation of Pbs2PP by unspecified phosphatase |
| k_pbs2P_dephosphorylation_mono | 1.08 | s^-1^ | Dephosphorylation of Pbs2P by unspecified phosphatase |

| **Pbs2 feedback** | |
| --- | --- |
| Pbs2 + Hog1PPc -> Pbs2_phosphorylated + Hog1PPc | 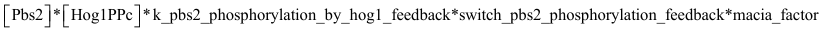 |
| Pbs2P + Hog1PPc -> Pbs2P_phosphorylated + Hog1PPc | 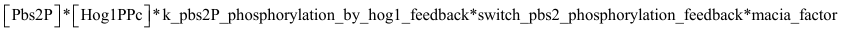 |
| Pbs2PP + Hog1PPc -> Pbs2PP_phosphorylated + Hog1PP | 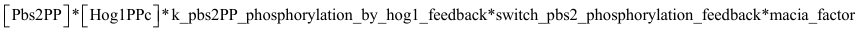 |
| Pbs2_phosphorylated -> Pbs2 | 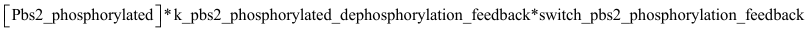 |
| Pbs2P_phosphorylated -> Pbs2P | 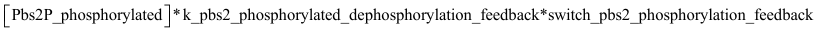 |
| Pbs2PP_phosphorylated -> Pbs2PP | 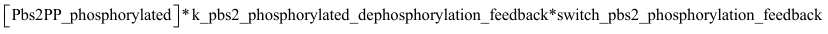 |
| Pbs2PP_phosphorylated -> Pbs2P_phosphorylated | 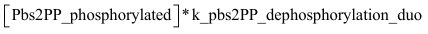 |
| Pbs2P_phosphorylated -> Pbs2_phosphorylated | 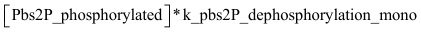 |

| Parameter | Parameter value | Unit | Additional Information |
| --- | --- | --- | --- |
| k_pbs2_phosphorylation_by_hog1_feedback | 6.07 | mM^-1^s^-1^ | Feedback phosphorylation of Pbs2 by double phosphorylated Hog1 |
| k_pbs2P_phosphorylation_by_hog1_feedback | 0.03 | mM^-1^s^-1^ | Feedback phosphorylation of Pbs2P by double phosphorylated Hog1 |
| k_pbs2PP_phosphorylation_by_hog1_feedback | 149.00 | mM^-1^s^-1^ | Feedback phosphorylation of Pbs2PP by double phosphorylated Hog1 |
| k_pbs2_phosphorylated_dephosphorylation_feedback | 0.38 | s^-1^ | Dephosphorylation of feedback phosphorylation by unspecified phosphatase |
| k_pbs2PP_dephosphorylation_duo | 0.83 | s^-1^ | Dephosphorylation of Pbs2PP by unspecified phosphatase |
| k_pbs2P_dephosphorylation_mono | 1.08 | s^-1^ | Dephosphorylation of Pbs2P by unspecified phosphatase |
| switch_pbs2_phosphorylation_feedback | 1 | au | Switch parameter to enable/disable pbs2 feedback |
| macia_factor | 1 | au | Factor that simulates inhibition of Hog1 activity when put to 0 |

| **Ssk2 feedback** | |
| --- | --- |
| Ssk2 + Hog1PPc -> Ssk2_phosphorylated + Hog1PPc | 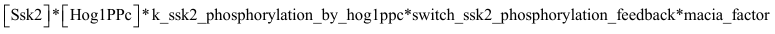 |
| Ssk2_phosphorylated + Hog1PPc -> Ssk2_phosphorylated_second + Hog1PPc | 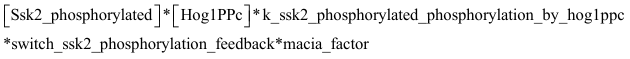 |
| Ssk2P + Hog1PPc -> Ssk2P_phosphorylated + Hog1PPc | 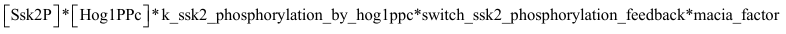 |
| Ssk2P_phosphorylated + Hog1PPc -> Ssk2P_phosphorylated_second + Hog1PPc | 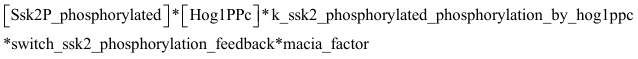 |
| Ssk2_phosphorylated_second -> Ssk2_phosphorylated | 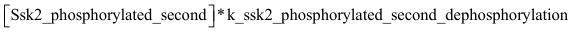 |
| Ssk2_phosphorylated -> Ssk2 | 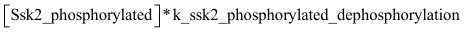 |
| Ssk2P_phosphorylated_second -> Ssk2P_phosphorylated | 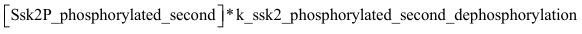 |
| Ssk2P_phosphorylated -> Ssk2P | 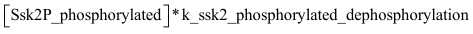 |
| Ssk2P_phosphorylated + Pbs2 -> Pbs2P + Ssk2P_phosphorylated | 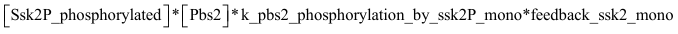 |
| Pbs2P + Ssk2P_phosphorylated -> Pbs2PP + Ssk2P_phosphorylated | 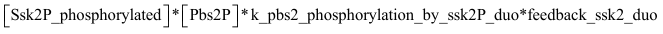 |
| Ssk2P_phosphorylated_second + Pbs2 -> Pbs2P + Ssk2P_phosphorylated_second | 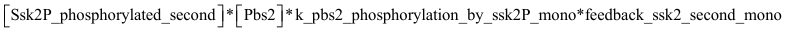 |
| Pbs2P + Ssk2P_phosphorylated_second -> Pbs2PP + Ssk2P_phosphorylated_second | 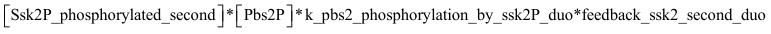 |
| Ssk2P_phosphorylated + Pbs2_phosphorylated -> Pbs2P_phosphorylated + Ssk2P_phosphorylated | 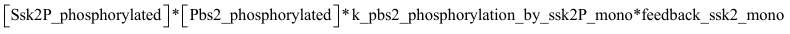 |
| Pbs2P_phosphorylated + Ssk2P_phosphorylated -> Pbs2PP_phosphorylated + Ssk2P_phosphorylated | 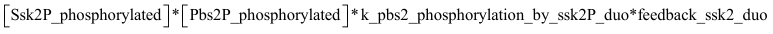 |
| Ssk2P_phosphorylated_second + Pbs2_phosphorylated -> Pbs2P_phosphorylated + Ssk2P_phosphorylated_second | 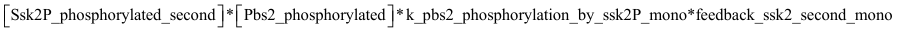 |
| Pbs2P_phosphorylated + Ssk2P_phosphorylated_second -> Pbs2PP_phosphorylated + Ssk2P_phosphorylated_second | 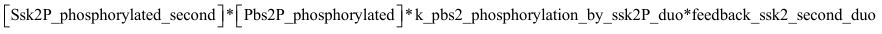 |
| **Unchanged reactions with Pbs2/Ssk2 feedback** | |
| Ssk2P + Pbs2_phosphorylated -> Pbs2P_phosphorylated + Ssk2P | 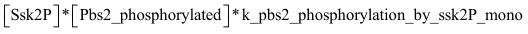 |
| Pbs2P_phosphorylated + Ssk2P -> Pbs2PP_phosphorylated + Ssk2P | 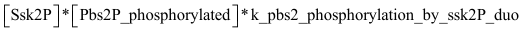 |

| Parameter | Parameter value | Unit | Additional Information |
| --- | --- | --- | --- |
| switch_ssk2_phosphorylation_feedback | 1 | au | Switch parameter to enable/disable feedback on Ssk2 |
| macia_factor | 1 | au | Factor that simulates inhibition of Hog1 activity when put to 0 |
| k_ssk2_phosphorylation_by_hog1ppc | 0.01 | mM^-1^s^-1^ | Feedback phosphorylation of Ssk2 by double phosphorylated Hog1 |
| k_ssk2_phosphorylated_phosphorylation_by_hog1ppc | 13.39 | mM^-1^s^-1^ | Feedback phosphorylation of phosphorylated Ssk2 by double phosphorylated Hog1 |
| k_ssk2_phosphorylated_second_dephosphorylation | 457.61 | s^-1^ | Dephosphorylation of feedback induced double phosphorylated Ssk2 by unspecified phosphatase |
| k_ssk2_phosphorylated_dephosphorylation | 5.02E-05 | s^-1^ | Dephosphorylation of feedback induced phosphorylated Ssk2 by unspecified phosphatase |
| k_pbs2_phosphorylation_by_ssk2P_mono | 448.75 | mM^-1^s^-1^ | Phosphorylation of Pbs2 by phosphorylated Ssk2 |
| k_pbs2p_phosphorylation_by_ssk2P_duo | 434.41 | mM^-1^s^-1^ | Phosphorylation of Pbs2P by phosphorylated Ssk2 |
| feedback_ssk2_mono | 1.92E-05 | mM^-1^s^-1^ | Multiplicative decrease of phosphorylation of Pbs2 by feedback induced monophosphorylated Ssk2 |
| feedback_ssk2_duo | 1.17E-05 | mM^-1^s^-1^ | Multiplicative decrease of phosphorylation of Pbs2P by feedback induced monophosphorylated Ssk2 |
| feedback_ssk2_second_mono | 0.84 | mM^-1^s^-1^ | Multiplicative decrease of phosphorylation of Pbs2 by feedback induced double phosphorylated Ssk2 |
| feedback_ssk2_second_duo | 7.85e-4 | mM^-1^s^-1^ | Multiplicative decrease of phosphorylation of Pbs2P by feedback induced double phosphorylated Ssk2 |

| Species | Initial value (unit) | Additional information |
| --- | --- | --- |
| Sln1_a_ | 0.0003 (mM) | Free Sln1 |
| Sln1P | 0.0407 (mM) | Free phosphorylated and activated Sln1 |
| Ypd1_b_ | 0.3926 (mM) | Free Ypd1 |
| Ypd1P | 0 (mM) | Free phosphorylated Ypd1 |
| Ssk1_c_ | 0 (mM) | Free Ssk1 activated |
| Ssk1P | 0.0744 (mM) | Free Ssk1 inactivated by phosphorylation |
| Ssk2_d_ | 0.0032 (mM) | Free Ssk2 inactive |
| Ssk2P | 0 (mM) | Free Ssk2 inactivated by phosphorylation |
| Ssk2_phosphorylated | 0.0084 (mM) | Free monophosphorylated, inactive Ssk2 |
| Ssk2_phosphorylated_second | 1.0720e-06 (mM) | Free double phosphorylated, inactive Ssk2 |
| Ssk2P_phosphorylated | 0.0019 (mM) | Free monophosphorylated, active Ssk2 |
| Ssk2P_phosphorylated_second | 1.0720e-06 (mM) | Free double phosphorylated, active Ssk2 |
| Pbs2_e_ | 0.1138 (mM) | Free Pbs2 |
| Pbs2P | 0.0029 (mM) | Free monophosphorylated Pbs2 |
| Pbs2PP | 0 (mM) | Free double activated Pbs2 |
| Pbs2_phosphorylated | 0.01 (mM) | Free Pbs2 with feedback phosphorylation |
| Pbs2P_phosphorylated | 3.4293e-05 (mM) | Free monophosphorylated Pbs2 with feedback phosphorylation |
| Pbs2PP_phosphorylated | 1.26e-6 (mM) | Free double activated Pbs2 with feedback phosphorylation |

_a_ Total amount of Sln1 in the cell: 656 molecules[1]

_b_ Total amount of Ypd1 in the cell: 6330 molecules[1]

_c_ Total amount of Ssk1 in the cell: 1200 molecules[1]

_d_ Total amount of Ssk2 in the cell: 217 molecules[1]

_e_ Total amount of Pbs2 in the cell: 2160 molecules[1]

1.	Ghaemmaghami, S., et al., *Global analysis of protein expression in yeast.* Nature, 2003. **425**(6959): p. 737-41.
