## Supplementary material for "Positive feedback induces switch between distributive and processive phosphorylation of Hog1": Supp table S7

| **Sho1 sub-branch** | |
| --- | --- |
| Reaction | Reaction rate |
| Sho1 -> Sho1_active | 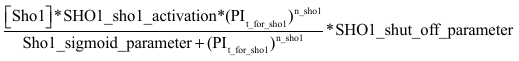 |
| Sho1_active -> Sho1 | 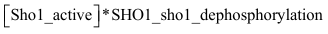 |
| Sho1 + Pbs2 -> Sho1_Pbs2 | 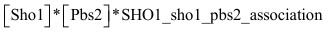 |
| Sho1_Pbs2 -> Sho1 + Pbs2 | 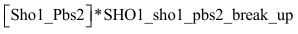 |
| Sho1_Pbs2 -> Sho1_active_Pbs2 | 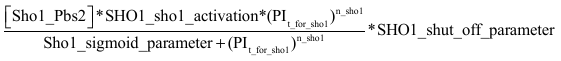 |
| Sho1_active + Pbs2 -> Sho1_active_Pbs2 | 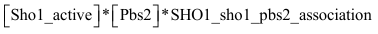 |
| Sho1_active_Pbs2 -> Sho1_active + Pbs2 | 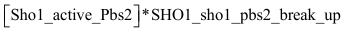 |
| 2 Ste11 + Ste50 -> Ste11_Ste50 | 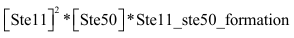 |
| Ste11_Ste50 -> 2 Ste11 + Ste50 | 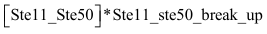 |
| Sho1_active_Pbs2 + Ste11_Ste50 -> Sho1_active_Ste11_Pbs2_inactive | 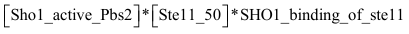 |
| Sho1_active_Ste11_Pbs2_inactive + Ste20 -> Sho1_active_Ste11_Pbs2 + Ste20 | 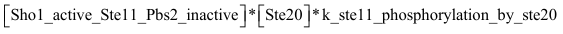 |
| Sho1_active + Ste11_Ste50 -> Sho1_active_Ste11 | 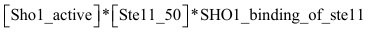 |
| Sho1_active_Ste11 + Ste20 -> Sho1_active_Ste11PP + Ste20 | 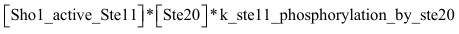 |
| Sho1_active_Ste11PP + Pbs2 -> Sho1_active_Ste11_Pbs2 | 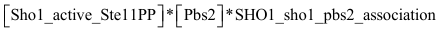 |
| Sho1_active_Ste11_Pbs2 -> Sho1_active_Ste11PP + Pbs2 | 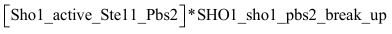 |
| Sho1_active_Ste11_Pbs2 -> Sho1_active_Ste11_Pbs2P | 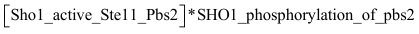 |
| Sho1_active_Ste11_Pbs2P -> Sho1_active_Ste11PP + Pbs2P | 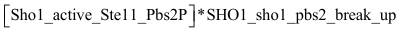 |
| Sho1_active_Ste11_Pbs2P -> Sho1_active_Ste11_Pbs2PP | 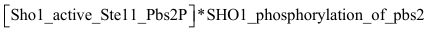 |
| Sho1_active_Ste11PP + Pbs2P -> Sho1_active_Ste11_Pbs2P | 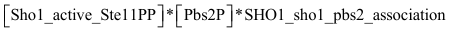 |
| Sho1_active_Ste11_Pbs2PP -> Sho1_active_Ste11PP + Pbs2PP | 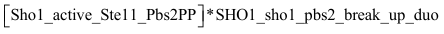 |
| Sho1_active_Ste11PP -> Sho1_active + Ste11P | 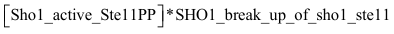 |
| Ste11PP -> Ste11_Ste50 | 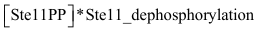 |

| Parameter | Parameter value | Unit | Additional information |
| --- | --- | --- | --- |
| SHO1_sho1_activation | 1.2221e+06 | au |  |
| n_sho1 | 9.7701 | au |  |
| PI_t_for_sho1_ | 1/PI_t_ |  | See **Volume Sub-model** for details of PI_t_ |
| Sho1_sigmoid_parameter | 1.1577e-04 | au |  |
| SHO1_shut_off_parameter | 1 | au | Switch parameter to enable/disable Sho1 sub-branch |
| SHO1_sho1_dephosphorylation | 0.0184 | s^-1^ | Dephosphorylation and thus deactivation of activated Sho1 |
| SHO1_sho1_pbs2_association | 7.7446 | mM^-1^s^-1^ | Association of Pbs2 to Sho1 |
| SHO1_sho1_pbs2_break_up | 0.5213 | s^-1^ | Disassociation of Pbs2 from Sho1 |
| Ste11_ste50_formation | 4.0738 | mM^-1^s^-1^ | Association of two Ste11 molecules and one Ste50 |
| Ste11_ste50_break_up | 0.8730 | s^-1^ | Disassociation of Ste11-Ste50 complex |
| SHO1_binding_of_ste11 | 1.4880e+06 | mM^-1^s^-1^ | Association of Sho1 and Ste11-Ste50 complex |
| k_ste11_phosphorylation_by_ste20 | 5.1928 | mM^-1^s^-1^ | Phosphorylation of Ste11 by Ste20 |
| SHO1_phosphorylation_of_pbs2 | 4.8295e+06 | s^-1^ | Phosphorylation of one Pbs2 phosphosite by activated Ste11 in big Sho1 complex |
| SHO1_sho1_pbs2_break_up_duo | 8.3426e+09 | s^-1^ | Disassociation of double phosphorylated Pbs2 and big Sho1 complex |
| SHO1_break_up_of_sho1_ste11 | 88.9406 | s^-1^ | Disassociation of Sho1 and Ste11-Ste50 complex |
| Ste11_dephosphorylation | 435.6122 | s^-1^ | Dephosphorylation of Ste11 by unspecified phosphatase |

| Species | Initial value (unit) | Additional information |
| --- | --- | --- |
| Sho1_a_ | 0.1334 (mM) | Free Sho1 |
| Sho1_active | 0 (mM) | Sho1 activated upon salt stress |
| Pbs2_b_ | 0.1138 (mM) | Free Pbs2 |
| Sho1_Pbs2 | 0.0102 (mM) | Sho1 bound to Pbs2 |
| Sho1_active_Pbs2 | 0 (mM) | Sho1 activated upon salt stress bound to Pbs2 |
| Ste11_c_ | 0.0421 (mM) | Free Ste11 |
| Ste50_d_ | 0.0263 (mM) | Free Ste50 |
| Ste11_Ste50 | 0 (mM) | Complex consisting of two molecules Ste11 and one Ste50 |
| Sho1_active_Ste11_Pbs2_inactive | 0 (mM) | Complex of activated Sho1, inactive Ste11-Ste50 complex and Pbs2 |
| Ste20_e_ | 0.0161 (mM) | Free Ste20 |
| Sho1_active_Ste11_Pbs2 | 0 (mM) | Complex of activated Sho1, activated Ste11-Ste50 complex and Pbs2 |
| Sho1_active_Ste11 | 0 (mM) | Complex of activated Sho1 and inactive Ste11-Ste50 complex |
| Sho1_active_Ste11PP | 0 (mM) | Complex of activated Sho1 and activated Ste11-Ste50 complex |
| Sho1_active_Ste11_Pbs2P | 0 (mM) | Complex of activated Sho1, activated Ste11-Ste50 complex and monophosphorylated Pbs2 |
| Pbs2P | 0.0029 (mM) | Free monophosphorylated Pbs2 |
| Sho1_active_Ste11_Pbs2PP | 0 (mM) | Complex of activated Sho1, activated Ste11-Ste50 complex and double phosphorylated Pbs2 |
| Pbs2PP | 0 (mM) | Free double phosphorylated Pbs2 |
| Ste11PP | 0 (mM) | Free activated Ste11-Ste50 complex |

_a_ Total amount of Sho1 in the cell: 2330 molecules[1]

_b_ Total amount of Pbs2 in the cell: 2160 molecules[1]

_c_ Total amount of Ste11 in the cell: 736 molecules[1]

_d_ Total amount of Ste50 in the cell: 1670 molecules[1]

_e_ Total amount of Ste20 in the cell: 259 molecules[1]

1.	Ghaemmaghami, S., et al., *Global analysis of protein expression in yeast.* Nature, 2003. **425**(6959): p. 737-41.
