## Supplementary information for "Positive feedback induces switch between distributive and processive phosphorylation of Hog1"

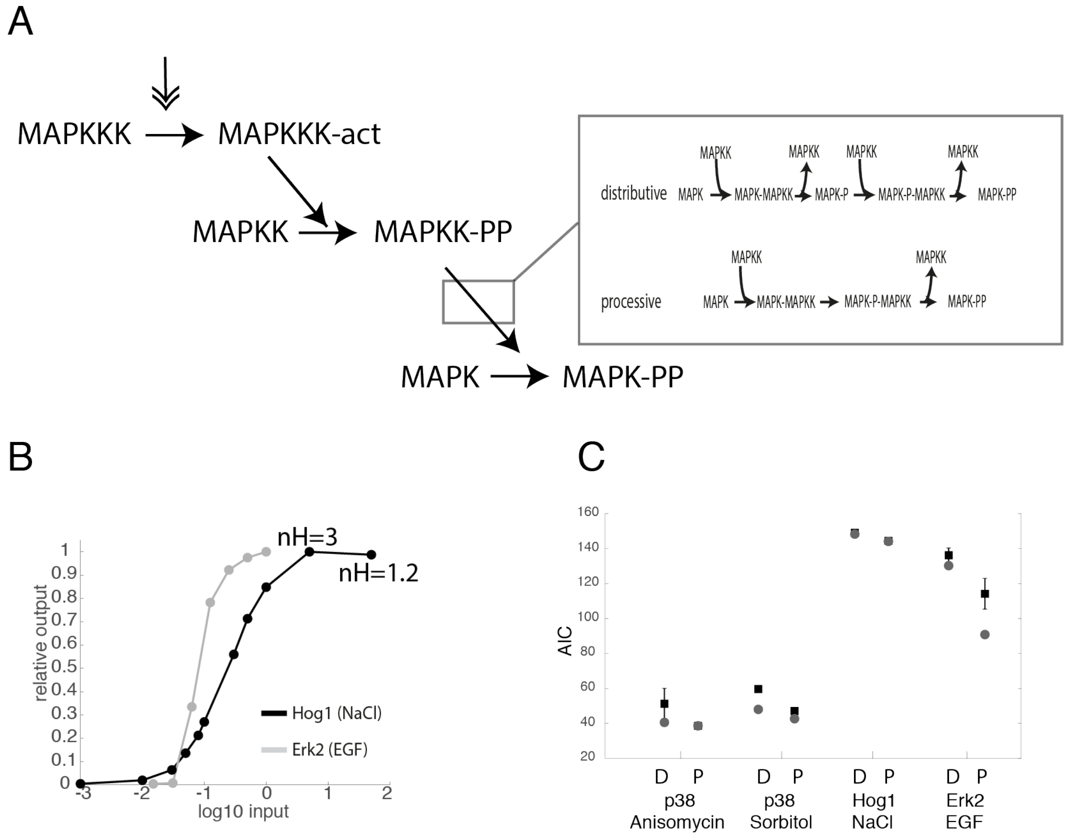


**Supp Figure S1: Modelling distributive or processive MAPK phosphorylation mechanisms**

(A) A schematic of the general MAPK model by (Huang and Ferrell, 1996). The two possible phosphorylation mechanisms, distributive or processive, are displayed. (B) Fitting a Hill function to data points of the activation of Hog1 by NaCl or Erk2 by EGF quantifies an ultrasensitive response of Hog1 activated by NaCl with a Hill coefficient of 3, and a graded response of Erk2 activated by EGF with a Hill coefficient 1.2. (C) A graph plotting the Akaike information Criterion (AIC) of the general MAPK model with either distributive (D) or processive (P) MAPK phosphorylation mechanism to different data sets. Mean of the best 100 fitting runs is displayed by a black square and the single best fit via a grey circle.

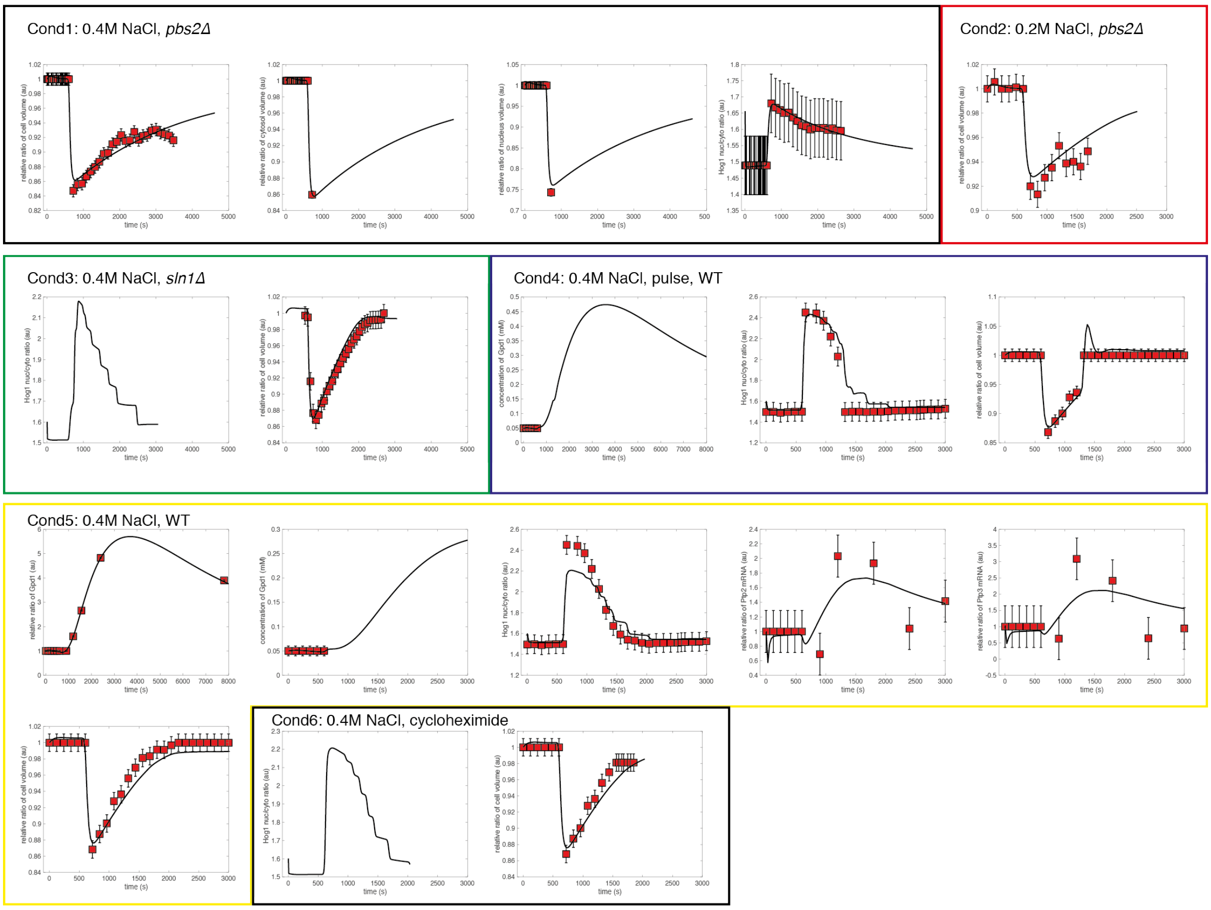


**Supp Figure S2: Data used for parameterization of the volume sub-model**

Data used for parameter optimization of the volume sub-model is shown as red squares. The simulation of the resulting best fit is indicated (solid black line). A total of six different conditions (boxes labeled Cond1-6), were fit simultaneously (see Supp Table S1 for details of data).

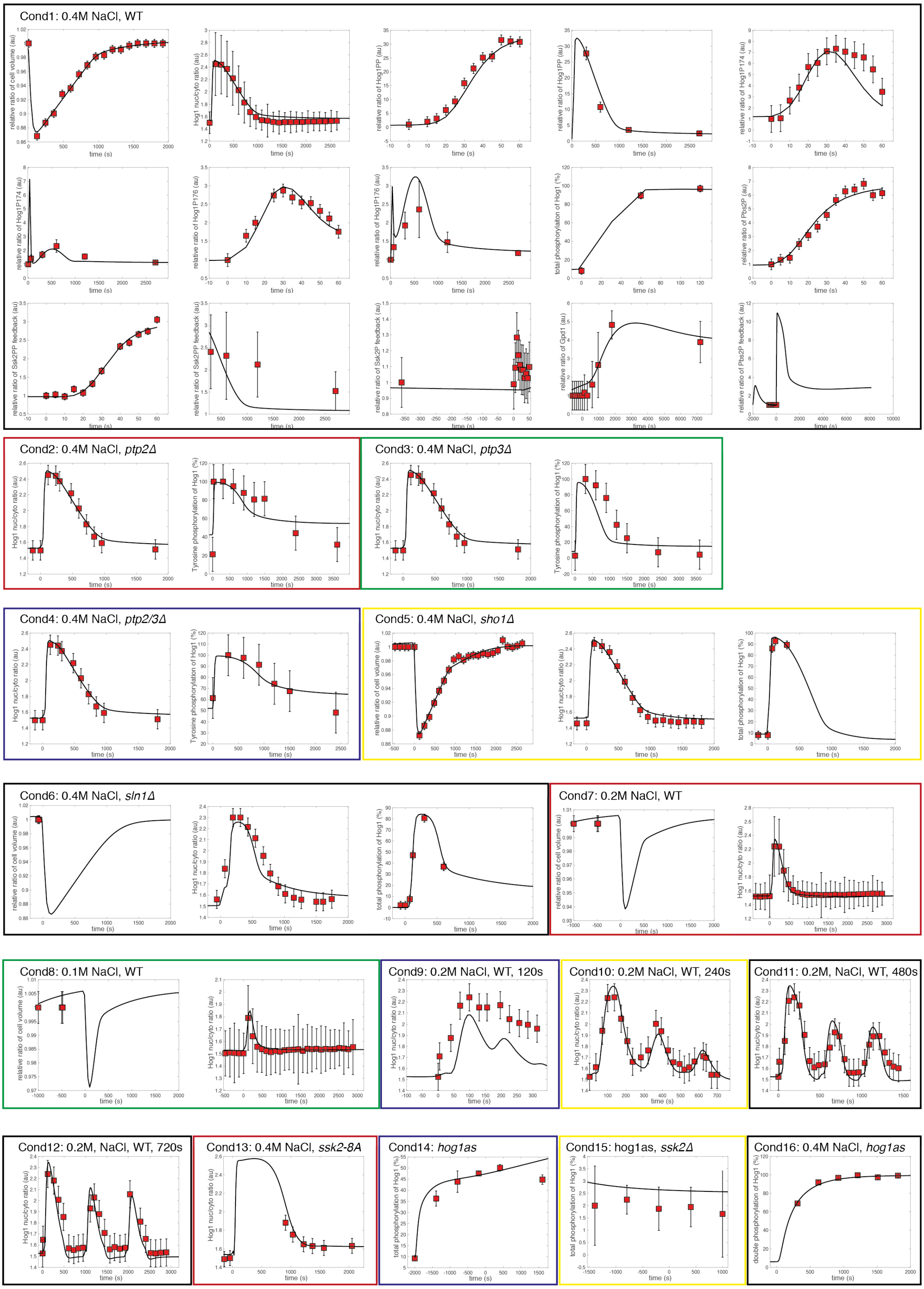


**Supp Figure S3: Data used for the parameterization of the overcomplete model**

Data points used for parameter optimization of the best fitting model as described in Figure 3B are shown as red squares. Note, that this data includes experimental measurements of monophosphorylated Hog1 species that were not previously used to generate the refined model (Figure 2B) that was used to distinguish between purely distributive and processive phosphorylation mechanisms and the presence or absence of feedback mechanisms. Thus, the best fitting model includes positive feedback on Pbs2, negative feedback, and a mixed phosphorylation mechanism as described in Figure 3A. Simulation of this model are indicated by the solid black line. The 16 conditions (boxes labeled Cond1-16) were fit simultaneously (see Supp Table S2 for details).

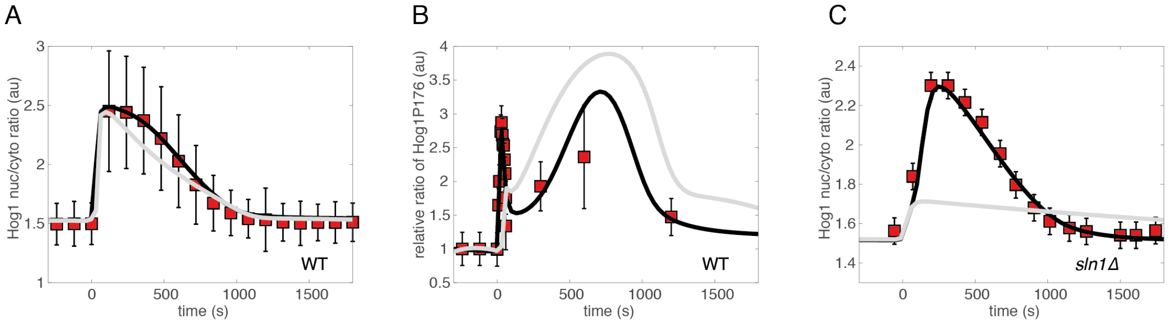


**Supp Figure S4: Differences between the best fitting results of the constrained and unconstrained processive model**

Data points used for parameter optimization of the full model are shown as red squares. Simulations of the resulting best fitting model from a topology with a processive phosphorylation mechanism and both negative feedback and positive feedback on Pbs2 are indicated for unconstrained (solid black line) and constrained (solid light grey) parameters. (A) Time courses of Hog1 nuclear to cytosolic ratio upon addition of 0.4M NaCl in wild type (WT) cells. (B) Time courses of the relative ratio between stimulated and basal levels of monophosphorylated Hog1-P176 upon addition of 0.4M NaCl in WT cells. (C) Time courses of Hog1 nuclear to cytosolic ratio upon addition of 0.4M NaCl in a *sln1Δ* strain.

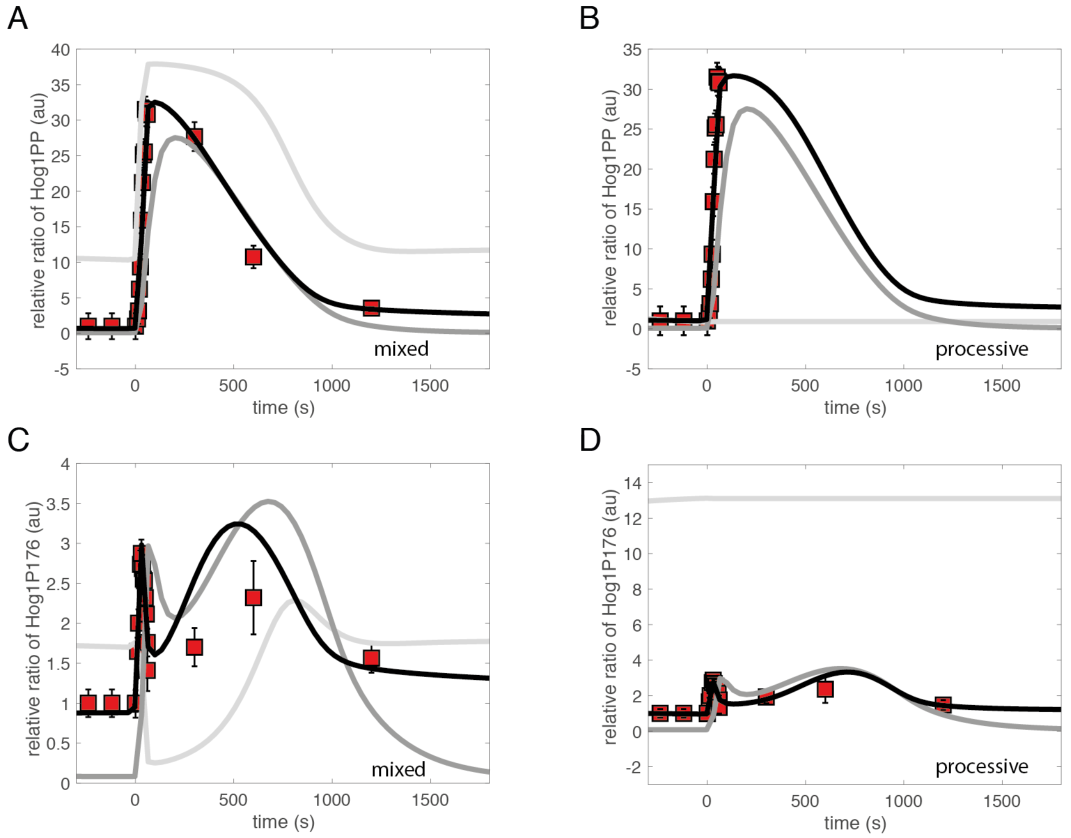


**Supp Figure S5: *In-silico* over- and under expression of Pbs2 in best fitting mixed or processive model**
(A-D) Simulations are shown of selected Hog1 species from the best fitting model with negative feedback on Ssk2, positive feedback on Pbs2 and either mixed or distributive phosphorylation mechanism of Hog1, optimized on data including the monophosphorylated Hog1 species. Graphs show experimental data points used for parameter optimization (red square) and simulations of the model with WT (solid black line), 0.1x (solid dark grey line) and 10x (solid light grey line) levels of Pbs2 concentrations. Time courses of the relative ratio between stimulated and basal levels following salt stimulation (0.4M NaCl) of dual phosphorylated Hog1-PP (A) and mono-phosphorylated Hog1-P176 (C) as the result of a mixed phosphorylation mechanism and dual phosphorylated Hog1-PP (B) and mono-phosphorylated Hog1-P176 (D) as the result of a processive phosphorylation mechanism are shown.

| Condition | Description | Source |
|---|---|---|
| Condition1 | 0.4M NaCl, *pbs2Δ* | this study |
| Condition2 | 0.2M NaCl, *pbs2Δ* | this study |
| Condition3 | 0.4M NaCl, *sln1Δ* | (Granados et al., 2017) |
| Condition4 | 0.4M NaCl, pulse | this study |
| Condition5 | 0.4M NaCl, WT | this study, (Petelenz-Kurdziel et al., 2013; Wurgler-Murphy et al., 1997) |
| Condition6 | 0.4M NaCl, Cycloheximide | (Mettetal et al., 2008) |

**Supp Table S1:**
Description of each condition and the data used for the parameter optimization of the volume sub-model. The best fit for each condition is described in Supp Figure S2.

| Condition | Description | Source |
|---|---|---|
| Condition1 | 0.4M NaCl, WT | this study,(English et al., 2015; Kanshin et al., 2015; Petelenz-Kurdziel et al., 2013; Sharifian et al., 2015; Vaga et al., 2014) |
| Condition2 | 0.4M NaCl, *ptp2Δ* | (Jacoby et al., 1997; Mattison and Ota, 2000) |
| Condition3 | 0.4M NaCl, *ptp3Δ* | (Jacoby et al., 1997; Mattison and Ota, 2000) |
| Condition4 | 0.4M NaCl, *ptp2/3Δ* | (Jacoby et al., 1997; Mattison and Ota, 2000) |
| Condition5 | 0.4M NaCl, *sho1Δ* | (Granados et al., 2017) |
| Condition6 | 0.4M NaCl, *sln1Δ* | (Granados et al., 2017) |
| Condition7 | 0.2M NaCl, WT | this study |
| Condition8 | 0.1M NaCl, WT | this study |
| Condition9 | 0.2M NaCl, pulse, t0=2min | (Mettetal et al., 2008) |
| Condition10 | 0.2M NaCl, pulse, t0=4min | (Mettetal et al., 2008) |
| Condition11 | 0.2M NaCl, pulse, t0=8min | (Mettetal et al., 2008) |
| Condition12 | 0.2M NaCl, pulse, t0=16min | (Mettetal et al., 2008) |
| Condition13 | 0.4M NaCl, *ssk2-8A* | (Sharifian et al., 2015) |
| Condition14 | *hog1as* | (Macia et al., 2009) |
| Condition15 | *hog1as, ssk2Δ* | (Macia et al., 2009) |
| Condition16 | 0.4M NaCl, *hog1as* | (English et al., 2015; Macia et al., 2009) |

**Supp Table S2:**
Description of each condition and the data used for the parameter optimization of the overcomplete model. The best fit for each condition is described in Supp Figure S3.

| Strain ID | Description | Background | Source | Used in figure |
| --- | --- | --- | --- | --- |
| BY4741 | MATa hisΔ1; leuΔ0; met15Δ0; ura3Δ0 | BY4741 | OpenBiosystems | Parental |
| W303 | leu2-3,112 trp1-1 can1-100 ura3-1 ade2-1 his3-11,15 | W303 | Lab collection | Parental |
| yMU49 | HTA2-CFP | BY4741 | Lab collection | Parental |
| yMM001 | HOG1-YFP HTA2-CFP | BY4741 | Lab collection | Parental |
| yMU19 | HOG1-YFP | W303 | Lab collection | Parental |
| yMM003 | HOG1-YFP HTA2-CFP Fus3_SKARS_reporter_mCherry_URA3 | BY4741 | Lab collection | Fig.2 Supp Fig.3 (Cond1) |
| yMM008 | Hog1-YFP HTA2-CFP Fus3_SKARS_reporter_mCherry_URA3 pbs2Δ | BY4741 | Lab collection | Supp Fig.2 (Cond1/2) |
| yMM004 | HOG1-YFP HTA2-CFP pSTL1-dPSTRr | BY4741 | Lab collection | Fig.2 Supp Fig.3 (Cond1) |
| yHS29 | HOG1-YFP HTA2-CFP pRPS2-Cherry-TMD | W303 | Lab collection | Fig.3C |

**Supp Table S3: Strain list**

| Plasmid ID | Description | Source |
|---|---|---|
| pED45 | Fus3_SKARS_reporter_mCherry_URA3 | (Durandau et al., 2015) |
| pSP135 | pAgTEF1-natMX-tAgTEF1 | Lab collection |
| pDA183 | pSIVU pRPL24A mCherry SZ2 tSIF2 -- pSTL1 UbiY 2xSv40NLS NewLinkerSZ1 | (Aymoz et al., 2016) |

**Supp Table S4: Plasmid list**
