## Supplementary material for "Positive feedback induces switch between distributive and processive phosphorylation of Hog1": Supp model files: arInputFunctionsC.c

```
#include <math.h>
#include "spline.c"
#include "monotone.c"

/* general input functions */
double heaviside(double t) {
    if (t < 0)
        return(0.0);
    else
        return(1.0);
}

/* delta functions are silently dropped */
double dirac(double t) {
    return 0.0;
}

double step1(double t, double u1, double t1, double u2) {
    if(t <= t1) {
        return(u1);
    } else {
        return(u2);
    }
}
double dstep1(double t, double u1, double t1, double u2, int p_index) {
    if(t <= t1) {
        if(p_index == 1)
            return(1.0);
        else
            return(0.0);
    } else {
        if(p_index == 3)
            return(1.0);
        else
            return(0.0);
    }
}

double step2(double t, double u1, double t1, double u2, double t2, double u3) {
    if(t <= t1) {
        return(u1);
    } else if( (t > t1) & (t <= t2) ) {
        return(u2);
    } else {
        return(u3);
    }
}
double dstep2(double t, double u1, double t1, double u2, double t2, double u3, int p_index) {
    if(t <= t1) {
        if(p_index == 1)
            return(1.0);
        else
            return(0);
    } else if( (t > t1) & (t <= t2) ) {
        if(p_index == 2)
            return(1.0);
        else
            return(0.0);
    } else {
        if(p_index == 3)
            return(1.0);
        else
            return(0.0);
    }
}

/* splines */

double spline3(double t, double t1, double p1, double t2, double p2, double t3, double p3, int ss, double dudt) {   
    double uout;
    
    double ts[3];
    double us[3];
    
    double b[3];
    double c[3];
    double d[3];
    
    ts[0] = t1;
    ts[1] = t2;
    ts[2] = t3;
    
    us[0] = p1;
    us[1] = p2;
    us[2] = p3;
    
    spline(3, ss, 0, dudt, 0.0, ts, us, b, c, d);
    uout = seval(3, t, ts, us, b, c, d);
    
    return(uout);
}

double monospline3(double t, double t1, double p1, double t2, double p2, double t3, double p3) {   
    double uout;
    
    double ts[3];
    double us[3];
    
    double b[3];
    double c[3];
    double d[3];
    
    ts[0] = t1;
    ts[1] = t2;
    ts[2] = t3;
    
    us[0] = p1;
    us[1] = p2;
    us[2] = p3;

    monotoneSpline( 3, ts, us, b, c, d );
    uout = seval(3, t, ts, us, b, c, d);
    
    return(uout);
}

double spline_pos3(double t, double t1, double p1, double t2, double p2, double t3, double p3, int ss, double dudt) {   
    int is;
    double uout;
    
    double ts[3];
    double us[3];
    double uslog[3];
    
    double b[3];
    double c[3];
    double d[3];
    
    ts[0] = t1;
    ts[1] = t2;
    ts[2] = t3;
    
    us[0] = p1;
    us[1] = p2;
    us[2] = p3;
    
    for (is = 0; is<3; is++){
        uslog[is] = log(us[is]);
    }
    
    spline(3, ss, 0, dudt, 0.0, ts, uslog, b, c, d);
    uout = seval(3, t, ts, uslog, b, c, d);
    
    return(exp(uout));
}

double spline4(double t, double t1, double p1, double t2, double p2, double t3, double p3, double t4, double p4, int ss, double dudt) {   
    double uout;
    
    double ts[4];
    double us[4];
    
    double b[4];
    double c[4];
    double d[4];
    
    ts[0] = t1;
    ts[1] = t2;
    ts[2] = t3;
    ts[3] = t4;
    
    us[0] = p1;
    us[1] = p2;
    us[2] = p3;
    us[3] = p4;
    
    spline(4, ss, 0, dudt, 0.0, ts, us, b, c, d);
    uout = seval(4, t, ts, us, b, c, d);
    
    return(uout);
}

double monospline4(double t, double t1, double p1, double t2, double p2, double t3, double p3, double t4, double p4) {   
    double uout;
    
    double ts[4];
    double us[4];
    
    double b[4];
    double c[4];
    double d[4];
    
    ts[0] = t1;
    ts[1] = t2;
    ts[2] = t3;
    ts[3] = t4;
    
    us[0] = p1;
    us[1] = p2;
    us[2] = p3;
    us[3] = p4;

    monotoneSpline( 4, ts, us, b, c, d );
    uout = seval(4, t, ts, us, b, c, d);
    
    return(uout);
}

double spline_pos4(double t, double t1, double p1, double t2, double p2, double t3, double p3, double t4, double p4, int ss, double dudt) {   
    int is;
    double uout;
    
    double ts[4];
    double us[4];
    double uslog[4];
    
    double b[4];
    double c[4];
    double d[4];
    
    ts[0] = t1;
    ts[1] = t2;
    ts[2] = t3;
    ts[3] = t4;
    
    us[0] = p1;
    us[1] = p2;
    us[2] = p3;
    us[3] = p4;
    
    for (is = 0; is<4; is++){
        uslog[is] = log(us[is]);
    }
    
    spline(4, ss, 0, dudt, 0.0, ts, uslog, b, c, d);
    uout = seval(4, t, ts, uslog, b, c, d);
    
    return(exp(uout));
}

double spline5(double t, double t1, double p1, double t2, double p2, double t3, double p3, double t4, double p4, double t5, double p5, int ss, double dudt) {   
    double uout;
    
    double ts[5];
    double us[5];
    
    double b[5];
    double c[5];
    double d[5];
    
    ts[0] = t1;
    ts[1] = t2;
    ts[2] = t3;
    ts[3] = t4;
    ts[4] = t5;
    
    us[0] = p1;
    us[1] = p2;
    us[2] = p3;
    us[3] = p4;
    us[4] = p5;
    
    spline(5, ss, 0, dudt, 0.0, ts, us, b, c, d);
    uout = seval(5, t, ts, us, b, c, d);
    
    return(uout);
}

double monospline5(double t, double t1, double p1, double t2, double p2, double t3, double p3, double t4, double p4, double t5, double p5) {   
    double uout;
    
    double ts[5];
    double us[5];
    
    double b[5];
    double c[5];
    double d[5];
    
    ts[0] = t1;
    ts[1] = t2;
    ts[2] = t3;
    ts[3] = t4;
    ts[4] = t5;
    
    us[0] = p1;
    us[1] = p2;
    us[2] = p3;
    us[3] = p4;
    us[4] = p5;

    monotoneSpline( 5, ts, us, b, c, d );
    uout = seval(5, t, ts, us, b, c, d);
    
    return(uout);
}

double spline_pos5(double t, double t1, double p1, double t2, double p2, double t3, double p3, double t4, double p4, double t5, double p5, int ss, double dudt) {   
    int is;
    double uout;
    
    double ts[5];
    double us[5];
    double uslog[5];
    
    double b[5];
    double c[5];
    double d[5];
    
    ts[0] = t1;
    ts[1] = t2;
    ts[2] = t3;
    ts[3] = t4;
    ts[4] = t5;
    
    us[0] = p1;
    us[1] = p2;
    us[2] = p3;
    us[3] = p4;
    us[4] = p5;
    
    for (is = 0; is<5; is++){
        uslog[is] = log(us[is]);
    }
    
    spline(5, ss, 0, dudt, 0.0, ts, uslog, b, c, d);
    uout = seval(5, t, ts, uslog, b, c, d);
    
    return(exp(uout));
}

double spline10(double t, double t1, double p1, double t2, double p2, double t3, double p3, double t4, double p4, double t5, double p5, double t6, double p6, double t7, double p7, double t8, double p8, double t9, double p9, double t10, double p10, int ss, double dudt) {   
    double uout;
    
    double ts[10];
    double us[10];
    
    double b[10];
    double c[10];
    double d[10];
    
    ts[0] = t1;
    ts[1] = t2;
    ts[2] = t3;
    ts[3] = t4;
    ts[4] = t5;
    ts[5] = t6;
    ts[6] = t7;
    ts[7] = t8;
    ts[8] = t9;
    ts[9] = t10;
    
    us[0] = p1;
    us[1] = p2;
    us[2] = p3;
    us[3] = p4;
    us[4] = p5;
    us[5] = p6;
    us[6] = p7;
    us[7] = p8;
    us[8] = p9;
    us[9] = p10;
    
    spline(10, ss, 0, dudt, 0.0, ts, us, b, c, d);
    uout = seval(10, t, ts, us, b, c, d);
    
    return(uout);
}

double monospline10(double t, double t1, double p1, double t2, double p2, double t3, double p3, double t4, double p4, double t5, double p5, double t6, double p6, double t7, double p7, double t8, double p8, double t9, double p9, double t10, double p10) {   
    double uout;
    
    double ts[10];
    double us[10];
    
    double b[10];
    double c[10];
    double d[10];
    
    ts[0] = t1;
    ts[1] = t2;
    ts[2] = t3;
    ts[3] = t4;
    ts[4] = t5;
    ts[5] = t6;
    ts[6] = t7;
    ts[7] = t8;
    ts[8] = t9;
    ts[9] = t10;
    
    us[0] = p1;
    us[1] = p2;
    us[2] = p3;
    us[3] = p4;
    us[4] = p5;
    us[5] = p6;
    us[6] = p7;
    us[7] = p8;
    us[8] = p9;
    us[9] = p10;
    
    monotoneSpline( 10, ts, us, b, c, d );
    uout = seval(10, t, ts, us, b, c, d);
    
    return(uout);
}

double spline_pos10(double t, double t1, double p1, double t2, double p2, double t3, double p3, double t4, double p4, double t5, double p5, double t6, double p6, double t7, double p7, double t8, double p8, double t9, double p9, double t10, double p10, int ss, double dudt) {   
    int is;
    double uout;
    
    double ts[10];
    double us[10];
    double uslog[10];
    
    double b[10];
    double c[10];
    double d[10];
    
    ts[0] = t1;
    ts[1] = t2;
    ts[2] = t3;
    ts[3] = t4;
    ts[4] = t5;
    ts[5] = t6;
    ts[6] = t7;
    ts[7] = t8;
    ts[8] = t9;
    ts[9] = t10;
    
    us[0] = p1;
    us[1] = p2;
    us[2] = p3;
    us[3] = p4;
    us[4] = p5;
    us[5] = p6;
    us[6] = p7;
    us[7] = p8;
    us[8] = p9;
    us[9] = p10; 
    
    for (is = 0; is<10; is++){
        uslog[is] = log(us[is]);
    }
    
    spline(10, ss, 0, dudt, 0.0, ts, uslog, b, c, d);
    uout = seval(10, t, ts, uslog, b, c, d);
    
    return(exp(uout));
}

/* spline derivatives */

double Dspline3(double t, double t1, double p1, double t2, double p2, double t3, double p3, int ss, double dudt, int id) {   
    double uout;
    
    double ts[3];
    double us[3];
    
    double b[3];
    double c[3];
    double d[3];
    
    ts[0] = t1;
    ts[1] = t2;
    ts[2] = t3;
    
    us[0] = 0.0;
    us[1] = 0.0;
    us[2] = 0.0;
    
    us[id-1] = 1.0;
    
    spline(3, ss, 0, dudt, 0.0, ts, us, b, c, d);
    uout = seval(3, t, ts, us, b, c, d);
    
    return(uout);
}

double Dmonospline3(double t, double t1, double p1, double t2, double p2, double t3, double p3, int id) {   
    double uout;
    
    double ts[3];
    double us[3];
    
    double b[3];
    double c[3];
    double d[3];
    
    ts[0] = t1;
    ts[1] = t2;
    ts[2] = t3;
    
    us[0] = 0.0;
    us[1] = 0.0;
    us[2] = 0.0;
    
    us[id-1] = 1.0;
    
    monotoneSpline( 3, ts, us, b, c, d );
    uout = seval(3, t, ts, us, b, c, d);
    
    return(uout);
}

double Dspline_pos3(double t, double t1, double p1, double t2, double p2, double t3, double p3, int ss, double dudt, int id) {   
    
    double uout;
    double uspline_pos3;
    double suspline3;
            
    double ps[3];
    
    ps[0] = p1;
    ps[1] = p2;
    ps[2] = p3;
    
    uspline_pos3 = spline_pos3(t, t1, p1, t2, p2, t3, p3, ss, dudt);
    suspline3 = Dspline3(t, t1, p1, t2, p2, t3, p3, ss, dudt, id);
    uout = suspline3 * uspline_pos3 * log(10.0);
    uout = uout / ps[id-1] / log(10.0);
    
    return(uout);
}

double Dspline4(double t, double t1, double p1, double t2, double p2, double t3, double p3, double t4, double p4, int ss, double dudt, int id) {   
    double uout;
    
    double ts[4];
    double us[4];
    
    double b[4];
    double c[4];
    double d[4];
    
    ts[0] = t1;
    ts[1] = t2;
    ts[2] = t3;
    ts[3] = t4;
    
    us[0] = 0.0;
    us[1] = 0.0;
    us[2] = 0.0;
    us[3] = 0.0;
    
    us[id-1] = 1.0;
    
    spline(4, ss, 0, dudt, 0.0, ts, us, b, c, d);
    uout = seval(4, t, ts, us, b, c, d);
    
    return(uout);
}

double Dmonospline4(double t, double t1, double p1, double t2, double p2, double t3, double p3, double t4, double p4, int id) {   
    double uout;
    
    double ts[4];
    double us[4];
    
    double b[4];
    double c[4];
    double d[4];
    
    ts[0] = t1;
    ts[1] = t2;
    ts[2] = t3;
    ts[3] = t4;
    
    us[0] = 0.0;
    us[1] = 0.0;
    us[2] = 0.0;
    us[3] = 0.0;
    
    us[id-1] = 1.0;
    
    monotoneSpline( 4, ts, us, b, c, d );
    uout = seval(4, t, ts, us, b, c, d);
    
    return(uout);
}

double Dspline_pos4(double t, double t1, double p1, double t2, double p2, double t3, double p3, double t4, double p4, int ss, double dudt, int id) {   
    
    double uout;
    double uspline_pos4;
    double suspline4;
            
    double ps[4];
    
    ps[0] = p1;
    ps[1] = p2;
    ps[2] = p3;
    ps[3] = p4;
    
    uspline_pos4 = spline_pos4(t, t1, p1, t2, p2, t3, p3, t4, p4, ss, dudt);
    suspline4 = Dspline4(t, t1, p1, t2, p2, t3, p3, t4, p4, ss, dudt, id);
    uout = suspline4 * uspline_pos4 * log(10.0);
    uout = uout / ps[id-1] / log(10.0);
    
    return(uout);
}

double Dspline5(double t, double t1, double p1, double t2, double p2, double t3, double p3, double t4, double p4, double t5, double p5, int ss, double dudt, int id) {   
    double uout;
    
    double ts[5];
    double us[5];
    
    double b[5];
    double c[5];
    double d[5];
    
    ts[0] = t1;
    ts[1] = t2;
    ts[2] = t3;
    ts[3] = t4;
    ts[4] = t5;
    
    us[0] = 0.0;
    us[1] = 0.0;
    us[2] = 0.0;
    us[3] = 0.0;
    us[4] = 0.0;
    
    us[id-1] = 1.0;
    
    spline(5, ss, 0, dudt, 0.0, ts, us, b, c, d);
    uout = seval(5, t, ts, us, b, c, d);
    
    return(uout);
}

double Dmonospline5(double t, double t1, double p1, double t2, double p2, double t3, double p3, double t4, double p4, double t5, double p5, int id) {   
    double uout;
    
    double ts[5];
    double us[5];
    
    double b[5];
    double c[5];
    double d[5];
    
    ts[0] = t1;
    ts[1] = t2;
    ts[2] = t3;
    ts[3] = t4;
    ts[4] = t5;
    
    us[0] = 0.0;
    us[1] = 0.0;
    us[2] = 0.0;
    us[3] = 0.0;
    us[4] = 0.0;
    
    us[id-1] = 1.0;
    
    monotoneSpline( 5, ts, us, b, c, d );
    uout = seval(5, t, ts, us, b, c, d);
    
    return(uout);
}

double Dspline_pos5(double t, double t1, double p1, double t2, double p2, double t3, double p3, double t4, double p4, double t5, double p5, int ss, double dudt, int id) {   
    
    double uout;
    double uspline_pos5;
    double suspline5;
            
    double ps[5];
    
    ps[0] = p1;
    ps[1] = p2;
    ps[2] = p3;
    ps[3] = p4;
    ps[4] = p5;
    
    uspline_pos5 = spline_pos5(t, t1, p1, t2, p2, t3, p3, t4, p4, t5, p5, ss, dudt);
    suspline5 = Dspline5(t, t1, p1, t2, p2, t3, p3, t4, p4, t5, p5, ss, dudt, id);
    uout = suspline5 * uspline_pos5 * log(10.0);
    uout = uout / ps[id-1] / log(10.0);
    
    return(uout);
}

double Dspline10(double t, double t1, double p1, double t2, double p2, double t3, double p3, double t4, double p4, double t5, double p5, double t6, double p6, double t7, double p7, double t8, double p8, double t9, double p9, double t10, double p10, int ss, double dudt, int id) {   
    double uout;
    
    double ts[10];
    double us[10];
    
    double b[10];
    double c[10];
    double d[10];
    
    ts[0] = t1;
    ts[1] = t2;
    ts[2] = t3;
    ts[3] = t4;
    ts[4] = t5;
    ts[5] = t6;
    ts[6] = t7;
    ts[7] = t8;
    ts[8] = t9;
    ts[9] = t10;
    
    us[0] = 0.0;
    us[1] = 0.0;
    us[2] = 0.0;
    us[3] = 0.0;
    us[4] = 0.0;
    us[5] = 0.0;
    us[6] = 0.0;
    us[7] = 0.0;
    us[8] = 0.0;
    us[9] = 0.0;
    
    us[id-1] = 1.0;
    
    spline(10, ss, 0, dudt, 0.0, ts, us, b, c, d);
    uout = seval(10, t, ts, us, b, c, d);
    
    return(uout);
}

double Dmonospline10(double t, double t1, double p1, double t2, double p2, double t3, double p3, double t4, double p4, double t5, double p5, double t6, double p6, double t7, double p7, double t8, double p8, double t9, double p9, double t10, double p10, int id) {   
    double uout;
    
    double ts[10];
    double us[10];
    
    double b[10];
    double c[10];
    double d[10];
    
    ts[0] = t1;
    ts[1] = t2;
    ts[2] = t3;
    ts[3] = t4;
    ts[4] = t5;
    ts[5] = t6;
    ts[6] = t7;
    ts[7] = t8;
    ts[8] = t9;
    ts[9] = t10;
    
    us[0] = 0.0;
    us[1] = 0.0;
    us[2] = 0.0;
    us[3] = 0.0;
    us[4] = 0.0;
    us[5] = 0.0;
    us[6] = 0.0;
    us[7] = 0.0;
    us[8] = 0.0;
    us[9] = 0.0;
    
    us[id-1] = 1.0;
    
    monotoneSpline( 10, ts, us, b, c, d );
    uout = seval(10, t, ts, us, b, c, d);
    
    return(uout);
}

double Dspline_pos10(double t, double t1, double p1, double t2, double p2, double t3, double p3, double t4, double p4, double t5, double p5, double t6, double p6, double t7, double p7, double t8, double p8, double t9, double p9, double t10, double p10, int ss, double dudt, int id) {   
    
    double uout;
    double uspline_pos10;
    double suspline10;
            
    double ps[10];
    
    ps[0] = p1;
    ps[1] = p2;
    ps[2] = p3;
    ps[3] = p4;
    ps[4] = p5;
    ps[5] = p6;
    ps[6] = p7;
    ps[7] = p8;
    ps[8] = p9;
    ps[9] = p10;
    
    uspline_pos10 = spline_pos10(t, t1, p1, t2, p2, t3, p3, t4, p4, t5, p5, t6, p6, t7, p7, t8, p8, t9, p9, t10, p10, ss, dudt);
    suspline10 = Dspline10(t, t1, p1, t2, p2, t3, p3, t4, p4, t5, p5, t6, p6, t7, p7, t8, p8, t9, p9, t10, p10, ss, dudt, id);
    uout = suspline10 * uspline_pos10 * log(10.0);
    uout = uout / ps[id-1] / log(10.0);
    
    return(uout);
}


    
/* custom rate laws */

double mmenten(double x, double vmax, double km){
    return(vmax * x / (km + x));
}

double mmenten_alt(double x, double klin, double ksat){
    return(0.0);
}

double hill_kd(double x, double h, double kd){
    return(pow(x,h) / (kd + pow(x,h)));
}

double hill_ka(double x, double h, double ka){
    /* return(pow(x,h) / (pow(ka,h) + pow(x,h))); */
    return(1 / (pow((ka/x),h) + 1));
}

/* try to get pressure function */
double PI_t_function(double V_os, double V_os_pi_t_0, double PI_t_0, double init_V_os, double output) {
    if(V_os > V_os_pi_t_0) {
        return((PI_t_0 * (V_os-V_os_pi_t_0))/((init_V_os-V_os_pi_t_0)));
    } else {
        return(output);
    }
}

double rebound_function(double V_os, double init_V_os, double output, double exponent, double normal_output) {
    if(V_os > init_V_os) {
        return((output * pow((V_os/init_V_os),exponent)));
    } else {
        return(normal_output);
    }
}

double test_function(double V_os, double V_os_pi_t_0, double output) {
    if(V_os > V_os_pi_t_0) {
        return(V_os - V_os_pi_t_0);
    } else {
        return(output);
    }
}
double divide_function(double V_os) {
        return(1/V_os);
   
}
```
