## Supplementary material for "Positive feedback induces switch between distributive and processive phosphorylation of Hog1": Supp model files: arInputFunctionsC.h

```
#include <math.h>

/* general input functions */
double heaviside(double t);
double dirac(double t);

double step1(double t, double u1, double t1, double u2);
double dstep1(double t, double u1, double t1, double u2, int p_index);

double step2(double t, double u1, double t1, double u2, double t2, double u3);
double dstep2(double t, double u1, double t1, double u2, double t2, double u3, int p_index);

/* splines */

double spline3(double t, double t1, double p1, double t2, double p2, double t3, double p3, int ss, double dudt);
double spline_pos3(double t, double t1, double p1, double t2, double p2, double t3, double p3, int ss, double dudt);
double monospline3(double t, double t1, double p1, double t2, double p2, double t3, double p3);

double spline4(double t, double t1, double p1, double t2, double p2, double t3, double p3, double t4, double p4, int ss, double dudt);
double spline_pos4(double t, double t1, double p1, double t2, double p2, double t3, double p3, double t4, double p4, int ss, double dudt);
double monospline4(double t, double t1, double p1, double t2, double p2, double t3, double p3, double t4, double p4);

double spline5(double t, double t1, double p1, double t2, double p2, double t3, double p3, double t4, double p4, double t5, double p5, int ss, double dudt);
double spline_pos5(double t, double t1, double p1, double t2, double p2, double t3, double p3, double t4, double p4, double t5, double p5, int ss, double dudt);
double monospline5(double t, double t1, double p1, double t2, double p2, double t3, double p3, double t4, double p4, double t5, double p5);

double spline10(double t, double t1, double p1, double t2, double p2, double t3, double p3, double t4, double p4, double t5, double p5, double t6, double p6, double t7, double p7, double t8, double p8, double t9, double p9, double t10, double p10, int ss, double dudt);
double spline_pos10(double t, double t1, double p1, double t2, double p2, double t3, double p3, double t4, double p4, double t5, double p5, double t6, double p6, double t7, double p7, double t8, double p8, double t9, double p9, double t10, double p10, int ss, double dudt);
double monospline10(double t, double t1, double p1, double t2, double p2, double t3, double p3, double t4, double p4, double t5, double p5, double t6, double p6, double t7, double p7, double t8, double p8, double t9, double p9, double t10, double p10);

/* spline derivatives */

double Dspline3(double t, double t1, double p1, double t2, double p2, double t3, double p3, int ss, double dudt, int id);
double Dspline_pos3(double t, double t1, double p1, double t2, double p2, double t3, double p3, int ss, double dudt, int id);
double Dmonospline3(double t, double t1, double p1, double t2, double p2, double t3, double p3, int id);

double Dspline4(double t, double t1, double p1, double t2, double p2, double t3, double p3, double t4, double p4, int ss, double dudt, int id);
double Dspline_pos4(double t, double t1, double p1, double t2, double p2, double t3, double p3, double t4, double p4, int ss, double dudt, int id);
double Dmonospline4(double t, double t1, double p1, double t2, double p2, double t3, double p3, double t4, double p4, int id);

double Dspline5(double t, double t1, double p1, double t2, double p2, double t3, double p3, double t4, double p4, double t5, double p5, int ss, double dudt, int id);
double Dspline_pos5(double t, double t1, double p1, double t2, double p2, double t3, double p3, double t4, double p4, double t5, double p5, int ss, double dudt, int id);
double Dmonospline5(double t, double t1, double p1, double t2, double p2, double t3, double p3, double t4, double p4, double t5, double p5, int id);

double Dspline10(double t, double t1, double p1, double t2, double p2, double t3, double p3, double t4, double p4, double t5, double p5, double t6, double p6, double t7, double p7, double t8, double p8, double t9, double p9, double t10, double p10, int ss, double dudt, int id);
double Dspline_pos10(double t, double t1, double p1, double t2, double p2, double t3, double p3, double t4, double p4, double t5, double p5, double t6, double p6, double t7, double p7, double t8, double p8, double t9, double p9, double t10, double p10, int ss, double dudt, int id);
double Dmonospline10(double t, double t1, double p1, double t2, double p2, double t3, double p3, double t4, double p4, double t5, double p5, double t6, double p6, double t7, double p7, double t8, double p8, double t9, double p9, double t10, double p10, int id);

/* custom rate laws */

double mmenten(double x, double vmax, double km);
double mmenten_alt(double x, double klin, double ksat);

double hill_kd(double x, double h, double kd);
double hill_ka(double x, double h, double ka);
double PI_t_function(double V_os, double V_os_pi_t_0, double PI_t_0, double init_V_os,double output);
double rebound_function(double V_os, double init_V_os,double output, double exponent, double normal_output);

double test_function(double V_os, double V_os_pi_t_0,double output);
double divide_function(double V_os);
```
